# MESMERISED: Super-accelerating T_1_ relaxometry and diffusion MRI with STEAM at 7 T for quantitative multi-contrast and diffusion imaging

**DOI:** 10.1101/2020.05.15.098269

**Authors:** F.J. Fritz, B.A. Poser, A. Roebroeck

## Abstract

There is an increasing interest in quantitative imaging of T_1_, T_2_ and diffusion contrast in the brain due to greater robustness against bias fields and artifacts, as well as better biophysical interpretability in terms of microstructure. However, acquisition time constraints are a challenge, particularly when multiple quantitative contrasts are desired and when extensive sampling of diffusion directions, high b-values or long diffusion times are needed for multi-compartment microstructure modeling. Although ultra-high fields of 7 T and above have desirable properties for many MR modalities, the shortening T_2_ and the high specific absorption rate (SAR) of inversion and refocusing pulses bring great challenges to quantitative T_1_, T_2_ and diffusion imaging. Here, we present the MESMERISED sequence (Multiplexed Echo Shifted Multiband Excited and Recalled Imaging of STEAM Encoded Diffusion). MESMERISED removes the dead time in Stimulated Echo Acquisition Mode (STEAM) imaging by an echo-shifting mechanism. The echo-shift (ES) factor is independent of multiband (MB) acceleration and allows for very high multiplicative (ESxMB) acceleration factors, particularly under moderate and long mixing times. This results in super-acceleration and high time efficiency at 7 T for quantitative T_1_ and diffusion imaging, while also retaining the capacity to perform quantitative T_2_ and B_1_ mapping. We demonstrate the super-acceleration of MESMERISED for whole-brain T_1_ relaxometry with total acceleration factors up to 36 at 1.8 mm isotropic resolution, and up to 54 at 1.25 mm resolution qT_1_ imaging, corresponding to a 6x and 9x speedup, respectively, compared to MB-only accelerated acquisitions. We then demonstrate highly efficient diffusion MRI with high b-values and long diffusion times in two separate cases. First, we show that super-accelerated multi-shell diffusion acquisitions with 370 whole-brain diffusion volumes over 8 b-value shells up to b = 7000 s/mm^2^ can be generated at 2 mm isotropic in under 8 minutes, a data rate of almost a volume per second, or at 1.8 mm isotropic in under 11 minutes, achieving up to 3.4x speedup compared to MB-only. A comparison of b = 7000 s/mm^2^ MESMERISED against standard MB pulsed gradient spin echo (PGSE) diffusion imaging shows 70% higher SNR efficiency and greater effectiveness in supporting complex diffusion signal modeling. Second, we demonstrate time-efficient sampling of different diffusion times with 1.8 mm isotropic diffusion data acquired at four diffusion times up to 290 ms, which supports both Diffusion Tensor Imaging (DTI) and Diffusion Kurtosis Imaging (DKI) at each diffusion time. Finally, we demonstrate how adding quantitative T_2_ and B_1_^+^ mapping to super-accelerated qT_1_ and diffusion imaging enables efficient quantitative multi-contrast mapping with the same MESMERISED sequence and the same readout train. MESMERISED extends possibilities to efficiently probe T_1_, T_2_ and diffusion contrast for multi-component modeling of tissue microstructure.

## Introduction

Magnetic resonance imaging (MRI) derives most of its versatility as an imaging technique from being able to weight images towards a multitude of contrasts, prominently among which are T_1_, T_2_/T_2_* and diffusion contrasts. Recently, there is an increasing interest in quantitative imaging of these contrasts for two main reasons. First, quantitative T_1_, T_2_ and diffusion images possess greater robustness against bias fields and artifacts, since the effects of spatially slowly varying radio frequency (RF) transmit and receive fields are mostly removed. Second, quantitative images have better biophysical interpretability, since they have interpretable physical units. This makes them a better proxy for biological microstructure and phenomena, such as myelin (e.g. Callaghan et al., 2015; Lutti et al., 2014; Weiskopf et al., 2013), g-ratio (e.g. Berman et al., 2019; Campbell et al., 2018), iron (e.g. Fukunaga et al., 2010; Stuber et al., 2014; Weiskopf et al., 2013), axonal density (e.g. Assaf and Basser, 2005; Zhang et al., 2012) as well as mesoscopic structure due to random axonal packing (e.g. Novikov et al., 2014). However, since quantitative MRI (qMRI) requires the acquisition of many volumes, from which the quantitative image is then derived, long acquisition times are a challenge, particularly when multiple quantitative contrasts are desired.

Acquisition time constraints are particularly pressing for diffusion MRI (dMRI). The signal varies with the orientation, strength and diffusion time encoded by diffusion gradients, making diffusion contrast inherently multi-dimensional. Moreover, recent developments in biophysical microstructure quantification with dMRI require sampling more of the multi-dimensional contrast space than with older methods. The conventional method for the analysis of white matter in dMRI imaging is the tensor model in Diffusion Tensor Imaging (DTI; Basser et al., 1994; Pierpaoli et al., 1996). Although a DTI derived index variable such as Fractional Anisotropy (FA) is sensitive, it is also unspecific, since differences in FA can reflect different axonal properties such as axon density, diameter distribution and myelination (Assaf et al., 2004; Assaf and Pasternak, 2008; Beaulieu, 2002; De Santis et al., 2014; Jones et al., 2013). Therefore, advances in diffusion microstructure modeling have aimed at greater specificity than DTI in relating the dMRI signal to the underlying cellular microstructure (Alexander et al., 2010; Assaf et al., 2013; Assaf and Basser, 2005; Assaf et al., 2008; Assaf et al., 2004; De Santis et al., 2014; Fieremans et al., 2013; Jelescu et al., 2015; Panagiotaki et al., 2012; Santis et al., 2014; Zhang et al., 2012). Models such as the Neurite Orientation Dispersion and Density Imaging model (NODDI; Zhang et al., 2012), the Combined Hindered And Restricted Model of Diffusion (CHARMED; Assaf and Basser, 2005), and the White Matter Tract Integrity model (WMTI; Fieremans et al., 2013) can provide measures specific to fiber density and orientation dispersion. These require the acquisition of a large number of diffusion directions over multiple b-values, sometimes extending to b-values as high as 6000 - 8000 s/mm^2^, easily leading to acquisition times of 10 - 20 minutes or longer. In addition, sensitivity can be gained to axonal diameter distributions (Alexander et al., 2010; Assaf et al., 2008; De Santis et al., 2016a; Zhang et al., 2011), water-exchange over cell membranes (Li et al., 2016; Nilsson et al., 2013) and mesoscopic structural effects due to random axonal packing (Burcaw et al., 2015; Novikov et al., 2014), but this requires the acquisition of a large number of b-values and diffusion times extending to 200 - 800 ms, often leading to acquisition times of 30 - 60 min.

Developments in echo-planar imaging (EPI) acquisition have allowed significant speed-up in the acquisition of volume time series weighted for T_1_, T_2_/T_2_* or diffusion. Prominently among these is multiband (MB) or simultaneous multi-slice (SMS) acquisition (Feinberg et al., 2010; Larkman et al., 2001; Moeller et al., 2010; Setsompop et al., 2012), which allows 2- to 4-fold speed-up for high flip angle applications at the expense of a higher specific absorption rate (SAR). However, even with MB acceleration, quantitative multi-contrast studies that require extensive diffusion MRI, as well as mapping of T_1_ and/or T_2_, are facing prohibitively long acquisition times.

In the domain of high spatial resolution imaging, ultra-high fields (UHF) of 7 Tesla (7 T) and above have enabled efficient high spatial resolution mapping of quantitative T_1_ (e.g. Marques et al., 2010; O’Brien et al., 2014), T_2_* (e.g. Fukunaga et al., 2010) and diffusion (e.g. Vu et al., 2015). In this work, our main aim was to establish whether the high signal-to-noise ratio (SNR) available at 7 T could be leveraged for highly time-efficient quantitative multi-contrast acquisition at moderate resolutions, particularly when including dMRI at high b-values or long diffusion times. Although UHF has desirable properties for many MR modalities, the shortening of T_2_ and the high SAR load of inversion and refocusing pulses bring challenges to efficient quantitative T_1_, T_2_ and diffusion imaging. Diffusion MRI sequences such as pulsed-gradient spin-echo (PGSE; e.g. Vu et al., 2015) and steady-state free precession (SSFP; e.g. Lu et al., 2012) have been optimized at 7 T to mitigate some of these limitations. However, they remain limited to MB acceleration factors of 2 to 4 (Eichner et al., 2014; Vu et al., 2015; Wu et al., 2013), with SAR as the main limiting factor.

Therefore, we investigated the possibilities of the Stimulated Echo Acquisition Mode (STEAM) sequence (Frahm et al., 1985). The STEAM sequence has some attractive advantages for multi-contrast MRI at UHF. First, it is highly multi-contrast capable, being able to provide quantitative T_1_ (qT_1_) contrast (by relaxometry on the mixing time, TM), quantitative T_2_ (qT_2_) contrast (by relaxometry on the echo time, TE), transmit (B_1_^+^) field mapping, and quantitative diffusion contrast. Furthermore, both b-value and diffusion time parameters in dMRI with STEAM can be probed at the high values desirable for many diffusion microstructure applications. Second, both a spin echo (SE) and stimulated echo (STE) signal can be independently acquired after a single excitation pulse, enhancing the capacity for efficient multi-contrast acquisition. Third, STEAM is highly efficient in SAR given its preferred 90°-90°-90° flip angle scheme. It has ∼60% of the SAR of a comparable 90°-180° spin-echo sequence and ∼33% of the SAR of a comparable 180°-90°-180° inversion-prepared spin-echo sequence (assuming similar slice-selection and a quadratic SAR/flip angle relationship). This also means that applying multiband pulses for SMS imaging is easier to apply for high MB factors in STEAM since SAR increases linearly with MB factor. However, the STEAM sequence also has inherent disadvantages. Both the SE and STE signals have only 50% of the signal level and signal to noise ratio (SNR), compared to the SE from a 90°-180° spin-echo sequence. Moreover, there is a considerable dead time in STEAM (Figure 1A), especially when moderate to long mixing times (useful for T_1_- weighting and probing of long diffusion times) are required, which leads to a low SNR per unit time efficiency.

**Figure 1:**
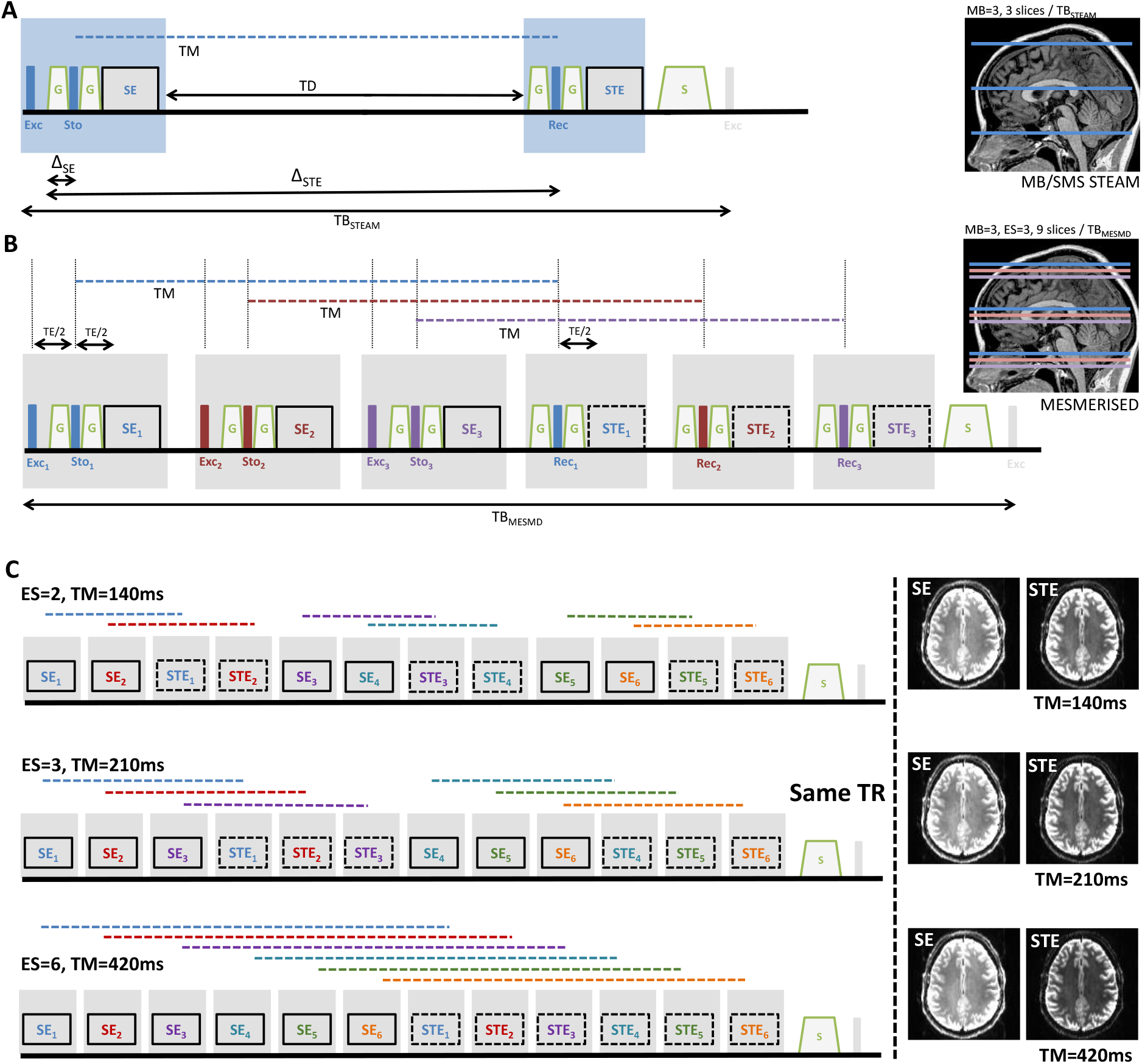
Multiband STEAM (MB-STEAM) versus MESMERISED. (A) A single block (TB_STEAM_) of the MB-STEAM sequence, consisting of a spin-echo (SE) and stimulated-echo (STE) readout, will acquire MB slices per block, leaving considerable dead time (TD), mostly determined by the desired mixing time (TM). (B) The MESMERISED sequence utilizes this dead time by echo-shifting one or more SE blocks (here ES factor 3), resulting in full duty cycle and high time efficiency. The stored longitudinal magnetization (horizontal dashed lines) of earlier SE blocks, evolves while later STE blocks are executed and SE_i_’s are acquired. Subsequently, STE_i_’s are read-out in the STE blocks. MESMERISED will acquire ESxMB slices per TB_MSMD_, which is slightly longer than TB_STEAM_. Exc/Sto/Rec: spatially selective Excitation/Storing/Recalling pulse; TE: echo time; G: diffusion/crusher gradient; S: spoiler gradient. (C) The block reordering property: MESMERISED enables a multitude of mixing times at the same TR with full duty cycle. Here SE and STE echoes are acquired for 6 slice groups, each containing a number of slices determined by the employed MB factor. SE and STE readout for the same slice groups, as well as the time evolution of longitudinally stored magnetization during the mixing time (dashed horizontal lines), are encoded by the same color. In MESMERISED each echo-shift factor (ES factor) engenders a simple reordering of the same sequence blocks to enable different mixing times at the same TR. This TR is the same for all allowable TM’s and is minimized by employing full acquisition duty cycle. The illustrated examples are for sequence block length of 70 ms, enabling TM = 140 ms for ES factor 2 (top), TM = 210 ms for ES = 3 (middle) and TM = 420 ms for ES = 6 (bottom).

Here, we present the MESMERISED sequence (Multiplexed Echo Shifted Multiband Excited and Recalled Imaging of STEAM Encoded Diffusion). MESMERISED removes the dead time in STEAM imaging by an echo-shifting mechanism. The echo-shift (ES) factor is independent of MB acceleration and allows for very high multiplicative ESxMB acceleration factors, particularly under moderate and long TM’s. This results in high duty cycle and high time efficiency at 7 T, particularly for quantitative T_1_ and diffusion imaging. In related work, Feinberg et al. (2010) introduced a different echo-shifting mechanism, termed simultaneous image refocusing (SIR) for gradient-echo EPI and pulsed-gradient spin echo (PGSE) EPI diffusion sequences. It also introduced the combined use of multiplexing SIR with MB acceleration to achieve very high multiplicative acceleration factors, closely related to the multiplexing of ES and MB in MESMERISED. Echo-shifting has been most applied in gradient-echo or steady-state free precession sequences to achieve echo times longer than their repetition time (Ehses et al., 2015; Gibson et al., 2006). Shifted slice orderings to make more effective use of dead time have been introduced early on in inversion recovery EPI for quantitative T_1_ mapping (Ordidge et al., 1990) and EPI based mapping has long been recognized for its high SNR per unit time (Mansfield et al., 1986). More recently, a multi-slice, inversion recovery EPI sequence (MS-IR-EPI) was proposed to enable T_1_ mapping with high spatial resolution and fidelity, making effective use of shifted slice orderings and simultaneous multi-slice acquisition (Panchuelo et al., 2021). The possibility of increasing the duty cycle of STEAM by exciting other slices or bands during the mixing time has recently been suggested (Zhang et al., 2019). To our knowledge, we here present the first demonstration of the echo-shifting mechanism in STEAM for quantitative T_1_ and diffusion mapping, and of the block reordering property (Figure 1C) which allows high acceleration and full acquisition duty cycle for multiple mixing times or diffusion times at the same TR.

We demonstrate the super-acceleration of MESMERISED for whole-brain T_1_ relaxometry with total acceleration factors up to 36 for 1.8 mm resolution and up to 54 for 1.25 mm resolution qT_1_ imaging. We then investigate highly efficient diffusion MRI with high b-values and long diffusion times. We show super-accelerated multi-shell diffusion acquisitions with 370 whole-brain diffusion volumes over 8 b-value shells up to b = 7000 s/mm^2^ at isotropic resolutions of 1.8 mm and 2.0 mm. We compare MESMERISED and standard MB PGSE for their SNR efficiency and their effectiveness in supporting complex diffusion signal modeling for the case of b-value = 7000 s/mm^2^ dMRI at 7 T. Subsequently, we demonstrate time-efficient sampling of different diffusion times with 1.8 mm isotropic diffusion data acquired at four diffusion times up to 290 ms. Finally, we demonstrate that quantitative T_2_ and B_1_^+^ mapping without echo-shifting acceleration can be added to achieve comprehensive multi-contrast mapping with the same MESMERISED sequence.

### Theory

The MESMERISED sequence is based on the standard (diffusion-weighted) STEAM sequence (Figure 1A). The STEAM pulse sequence uses three radiofrequency (RF) pulses: excitation (Exc), storing (Sto) and recalling (Rec) pulses, which can create several spin echoes (SEs) and a stimulated echo (STE), which was first recognized by Erwin Hahn (Hahn, 1950) and later introduced to MRI by Frahm and colleagues (Frahm et al., 1985). In general terms, the excitation pulse places the magnetization into the transverse plane where it undergoes T_2_^*^ decay. Subsequently, the storing pulse places (‘stores’) part of the total magnetization along the longitudinal axis where it is subject to T_1_ decay during the mixing time (TM). The remaining part of the magnetization forms the primary SE signal, also known as an “eight-ball echo” (Hahn, 1950) or Hahn echo, since it is the lesser known form of a spin echo formed by a 90°-90° pulse pair. Finally, the recalling pulse places (‘recalls’) the stored magnetization to the transverse plane where it once again undergoes T_2_^*^ decay until the STE is formed. This sequence of pulses can create three more spin-echo signals (the second, third and fourth spin-echo) after the STE, which are the consequence of the free induction decay (FID) signal generated by the storing and recalling pulses. We focus on the first or primary SE signal in the remainder of this work and refer to it simply as ‘the SE’ hereafter. Whereas the SE undergoes only T_2_ decay, the STE signal undergoes both T_1_ and T_2_ decay, because the signal is obtained from spins *stored* along the longitudinal axis for the duration of the TM, during which there is T_1_ relaxation. Furthermore, both echoes (SE and STE) share the transverse magnetization created by the excitation pulse and therefore split the total excited signal, which becomes evenly distributed between SE and STE when the storing pulse has a flip angle of 90°.

MESMERISED removes the dead time in STEAM by shifting Exc-Sto pulse pairs and SE readout for one or more MB slice groups into the dead time of earlier slice groups (Figure 1B). We refer to this as echo-shifting, by analogy to sequences that shift sequence elements to achieve echo times longer than their repetition time (TR) in gradient-echo or steady-state free precession sequences (c.f. Ehses et al., 2015; Gibson et al., 2006). However, we note the shifting mechanism here is fundamentally different and does not achieve TE > TR. The echo-shifting in MESMERISED temporally interleaves and multiplexes the contrast evolution of several slices or MB slice-groups. The stored longitudinal magnetization of earlier Exc-Sto and SE readout sequence blocks (SE_i_ blocks) evolves while later SE_i+n_ blocks are executed and SEs are acquired. Subsequently, STEs are recalled and read-out in a sequence of STE_i_ blocks (Figure 1B), placed at time TM to their corresponding SE_i_ blocks. We refer to the number of echo-shifted slice groups as the echo-shift factor (ES factor). The echo-shifting is independent of MB acceleration and allows for very high multiplicative ESxMB acceleration factors, particularly under moderate and long TM’s. This results in full duty cycle and high time efficiency, particularly for quantitative T_1_ and diffusion imaging.

For instance, T_1_ relaxometry with STEAM requires a range of TM’s acquired under the same TR and TE. Whereas total acquisition time of T_1_ relaxometry with MB-STEAM would be determined by the TR of the *longest* TM, acquisition time of T_1_ relaxometry with MESMERISED is limited by the TR of *shortest* possible TM. Here, MESMERISED makes use of its *block reordering property* (Figure 1C): a different choice of ES factor determines a simple reordering of the same sequence blocks and this enables a different mixing time to be acquired at the same minimized TR. Therefore, in MESMERISED all TM’s or diffusion times required, for instance to accurately fit an exponential decay model, can be acquired at full acquisition duty cycle and the same short TR.

For diffusion imaging, mono-polar diffusion gradient pairs are inserted symmetrically around the storing and recalling pulses. Although only the first (before the storing pulse) and fourth (after the recalling pulse) gradients are needed for diffusion weighting of the STE signal, there are several advantages to a double time-symmetric placement of four gradients. First, these also serve as crusher gradients at low amplitude and duration in non-diffusion-weighted imaging, helping to remove unwanted echo pathways. Second, the first pair of gradients create a pulsed-gradient spin echo (PGSE) acquisition in the SE_i_ block, with the corresponding PGSE diffusion contrast in the SE. Third, the second and third gradient act as spoilers during the mixing time for unwanted (secondary and tertiary) spin echo pathways (as shown in a phase graph, see Supplementary Figure 1). This spoiler action is enhanced in MESMERISED, since the gradients of echo-shifted blocks add to the total spoiler-moment during TM. Additionally, ‘unbalancing’ the third gradient with respect to the fourth by modifying its moment provides a further degree of adjustment to avoid unwanted echo pathways (cf. Supplementary Figure 1). MESMERISED can acquire diffusion-weighted signals at long diffusion times and high b-values in its STE signal, at relatively low duty cycle demands on the gradients. In addition, it allows highly accelerated probing of multiple diffusion times at the same TR and TE using block reordering property (Figure 1C) in the same way as in T_1_ relaxometry. Finally, as a STEAM variant with both a SE and STE readout, MESMERISED can acquire both a low b-value and short diffusion time (in its diffusion-weighted SEs) and a high b-value and long diffusion time (in its diffusion-weighted STEs) in a single TR.

## Methods

### MESMERISED sequence implementation

The MESMERISED sequence was implemented on a Siemens 7 T MAGNETOM platform (Siemens Healthineers Erlangen, Germany). A sequence unit (or building block) was created consisting of a single multiband selective pulse surrounded by two gradient pulses and followed by an echo-planar imaging (EPI) readout. This unit serves both as an STE block and, when including a matched multiband pulse at the outset, as an SE block (Figure 1B). Blipped-CAIPIRINHA simultaneous multi-slice reconstruction (Setsompop et al., 2012) with the split-slice GRAPPA leak-block approach (Cauley et al., 2014) was performed by using the online reconstruction that is part of the MGH SMS-EPI release for Siemens platforms (https://www.nmr.mgh.harvard.edu/software/c2p/sms). In-plane acceleration was reconstructed with GRAPPA (Griswold et al., 2002) using FLEET autocalibration scans as described in (Blazejewska et al., 2017). The MESMERISED implementation allows for user-controllable MB, GRAPPA and ES acceleration factors, along with the common parameters (TE, TR, number of slices, and resolution parameters: slice thickness, in-plane resolution, readout bandwidth). For diffusion imaging, duration δ, amplitude |G| and time spacing Δ of the diffusion gradient pairs are user-controllable to determine b-value and diffusion time for the SE (b_SE_ and Δ_SE_= Δ) and the STE (b_STE_ and Δ_STE_= Δ+TM). Diffusion gradient direction tables were implemented as multiples of 12 directions (24, 36, 48, 60) distributed around a full sphere by an electrostatic repulsion algorithm (Jones, 2004), interspersed with b_0_ volumes every 12 directions. Multi-shell tables were implemented by combining tables and suitable modulation of |G| by scaling the length of the diffusion direction vector. To remove the need for SAR-intensive fat saturation (FatSat) pulses, “fat-unfocusing” as described in (Ivanov et al., 2010) was achieved by the appropriate setting of the pulse lengths and bandwidth-time product (BWTP).

### MRI data acquisition

Whole-brain acquisitions were performed on a whole-body research 7 T Siemens MAGNETOM (Siemens Healthcare, Erlangen, Germany) system with gradients delivering 70 mT/m maximum strength at a maximum slew rate of 200 T/m/s per physical axis. For RF transmission and reception, a single-channel-transmit/32-channel-receive head coil (Nova Medical, Wilmington, MA, USA) was used. Written informed consent was obtained from all the subjects before imaging and all procedures were approved by the Ethics Review Committee Psychology and Neuroscience (ERCPN) of the faculty of Psychology and Neuroscience, Maastricht University. MESMERISED was tested on six healthy subjects (4 males and 2 females, average weight 75 ±+-5 kg and age 29 ±+-3 years). Dielectric pads (Teeuwisse et al., 2012) were used to improve B_1_^+^ field homogeneity by positioning two pads at the ears/temples. All sessions started with a localizer scan in three orthogonal directions. Subsequently, a B_0_ field map was acquired (Cusack and Papadakis, 2002) and used to optimized B_0_ shim currents. An absolute B_1_^+^ map (using a pre-saturation turbo-flash (PreSat-TFL) sequence; Chung et al., 2010) was then acquired and used to calibrate the global transmit flip angle.

Subsequently, MESMERISED acquisitions were performed with full brain coverage, with slices in a transverse orientation. All diffusion and crusher gradients were run in full symmetric format, i.e. unbalancing of the third gradient was not needed to avoid echo pathway contamination in the experiments reported here. EPI readout was executed along the anterior-posterior (AP) phase-encoded direction, and undersampled with GRAPPA acceleration factor 2 or 3 and using a partial Fourier factor of 6/8. In-plane reference (ACS) lines for GRAPPA (24 lines for factor 2, 36 lines for factor 3) were acquired using FLEET at a flip angle 12° with 2 to 5 dummy pre-scans. For blipped CAIPI MB (Breuer et al., 2006; Setsompop et al., 2012), a CAIPI-shift (CS) factor was employed (MB: 3, CS: 2; and MB: 4, CS: 3). Every acquisition was accompanied by 5 volumes with a reversed-phase encoding direction (posterior-anterior) only for the contrast with highest signal (i.e. shortest TM or TE, flip angles 90°-90°-90°, and/or b_0_ acquisition), used for correction of off-resonance distortions. For all the acquisitions, the slices were acquired in an interleaved ordering without slice gap.

MESMERISED protocols were created for imaging of quantitative T_1_ (qT_1_), quantitative T_2_ (qT_2_), transmit field (B_1_^+^) mapping and diffusion. In MESMERISED the number of slices must be an integer multiple of ESxMB for all employed ES and MB factors. For qT_2_, B_1_^+^ and diffusion protocols with a single mixing time or diffusion time this provides a single constraint. For qT_1_ or diffusion protocols with multiple TM or diffusion times (Δ), this provides a set of constraints, one for each ES/TM combination. These constraints are easily solved by choosing slice numbers N with a large number of integer divisors (highly composite numbers), such as 60, 72, 90 and 108, which can be used for isotropic imaging resolutions of 2 mm, 1.8 mm, 1.5 mm, and 1.25 mm, respectively. Each integer divisor of slice number divided by MB factor (i.e. N/MB), then determines an allowable ES factor under the same TR (because it determines a single temporal reordering of sequence blocks, see Figure 1C). For instance, at 72 slices with MB = 3, N/MB = 24 the allowable ES factors are [1, 2, 3, 4, 6, 8, 12, 24]. At full duty cycle, each such ES is associated with a single minimum TM, determined by the length of RF pulses, gradient pulses, and EPI train length. Again due to MESMERISED’s temporal block reordering property, the minimum TM for each allowable ES is (very close to) ES*TM_ES=1_, i.e. an integer multiple of the minimum TM for ES = 1.

A proof of principle test was performed to demonstrate that MESMERISED’s echo-shifting does not affect the resulting SE and STE images, and therefore the same signal can be acquired in shorter total acquisition time. Sets of MESMERISED SE and STE images were acquired at 1.8 mm isotropic resolution at different ES but fixed TM = 300 ms, TR = 11.88 s and TE = 47.5 ms. ESxMB combinations of 1×3, 2×3 and 4×3 were acquired with transverse slices, GRAPPA 2, FoV 230 × 230 mm^2^ and partial Fourier 6/8. The achievable (but unused) minimum TR for ESxMB= 2×3 and 4×3 were recorded to quantify the acceleration potential of MESMERISED.

### qT_1_ acquisition

Quantitative T_1_ imaging was performed at 1.8 mm isotropic and 1.25 mm isotropic, by performing relaxometry over a range of allowable ES/TM combinations at the same TR and TE. Five repetitions of each TM were acquired to assess retrospectively what level of data averaging (1, 3 or 5 repetitions) supports robust exponential T_1_-decay model fitting. Additionally, T_1_ relaxometry was performed both at the minimum possible TR, at full duty cycle and acceleration, and with a doubling of TR and all TM’s. This was done to assess the range of TM’s necessary for robust exponential T_1_- decay model fitting. Full acquisition parameters are given in Supplementary Table 1.

### Diffusion acquisition

Multi-shell diffusion imaging was performed at 1.8 mm and 2.0 mm isotropic for different maximum b-values. Diffusion acquisitions were designed to explore the strengths of MESMERISED in achieving a high number of volumes per unit time, and either high b-values or a large range of Δ’s. A set of multi-shell diffusion acquisitions with maximum b-value 7000 s/mm^2^, 185 directions over 4 shells (a total of 370 SE and STE volumes), were performed at 1.8 mm isotropic and different ES/TM, realizing different diffusion times between 80 ms and 430 ms in the STE. A 4-shell acquisition at 2 mm isotropic with maximum b-value 7000 s/mm^2^ was performed to explore the effect of a very short TR (2.36 sec) and a very high number of volumes per unit time.

MESMERISED and standard MB PGSE were compared for their signal-to-noise efficiency in b-value = 7000 s/mm^2^ dMRI at 7 T and their effectiveness in supporting complex diffusion signal modeling. In order to achieve an objective comparison under the same RF pulse, EPI readout and image reconstruction implementations, the same MESMERISED-capable sequence was run in PGSE mode by omitting the STE block and modifying the storing pulse flip angle (180°) and phase. Whole brain (72 slices) 1.8 mm isotropic b = 7000 s/mm^2^ diffusion-weighted MESMERISED with 4 echo-shift factors (ESxMB: 2×3, 3×3, 4×3 and 6×3; TE: 49.9, 46.2, 43.8, 40.9 ms; TM: 140, 210, 280 and 420 ms; SE b-val: 752, 483, 350, 223 s/mm^2^, respectively) and PGSE (MB:2; TR: 4.21 s.; TE: 97.8 ms) were optimized to run just within SAR and gradient duty cycle limits. For each, 60 directions at b = 7000 s/mm^2^ with 6 b_0_s (for diffusion microstructure modelling) as well as 27 matched b_0_s (for temporal SNR and SNR efficiency calculation) were acquired in one healthy subject (male, 79 kg). MESMERISED at MB = 3 and PGSE at MB = 2 were selected, since under basic assumptions (SAR grows linearly with MB factor and quadratic with flip angle) these settings would have a similar SAR load (within 10% at the same TR with MESMERISED-MB3 10% more efficient than PGSE-MB2). Pulse lengths (4.48 ms for excitation, 8.96 ms for refocusing, storing and recalling) were matched between MESMERISED and PGSE and chosen to satisfy “fat-unfocusing” constraints as described in (Ivanov et al., 2010), as well as balancing a reasonable length (compared to white matter T_2_*) with SAR constraints. A potential TE shortening advantage for MESMERISED by instead running the excitation pulse as the long pulse and storing and recalling as the short pulse was avoided to keep the RF-pulse regime as similar as possible. Flip angle calibration was performed as previously described for 7 T diffusion acquisition (Vu et al., 2015), which lead to a reference voltage of 235 V for the 79 kg subject and ∼15-20% overflipping in the center of the brain. SAR constraints alone forced PGSE to run at a TR of close to 4 s, far higher than the full acquisition duty cycle TR of ∼2.5 s. Moreover, gradient duty cycle constraints forced further TR compromises for PGSE as established by running the protocol with the evenly sphere-sampled 60 direction table at the desired b = 7000 s/mm^2^ while adjusting diffusion gradient amplitude and duration (and by extension TE and TR) until the full protocol ran without gradient power amplifier overloads. Gradient limits forced b = 7000 s/mm^2^ PGSE to run at |G|: 49 mT/m, lengthening TR and TE (to 4.21 s. and 97.8 ms, respectively), whereas for MESMERISED this allowed full diffusion gradient amplitude (|G|: 70 mT/m) and full acquisition duty cycle TR (3.4 s). EPI readout (GRAPPA-2 with FLEET ACS, FoV 218 × 218 mm^2^ and PF 6/8) was matched for echo-spacing (0.57 ms) and for readout bandwidth within 5% (MESMERISED: 2164 Hz/pix; PGSE: 2066 Hz/pix). Forcing a full EPI bandwidth match would increase PGSE TE and TR further through a peripheral nerve stimulation limit that did not occur in MESMERISED at their respective TRs. All four MESMERISED protocols and the PGSE protocol ran between 90% and 100% of the 6-minute SAR limit.

Finally, to assess efficient exploration of diffusion-time dependent effects on mean diffusivity and mean kurtosis with MESMERISED, an acquisition at 1.8 mm isotropic at four different diffusion times and two different b-values (1000 and 2500 s/mm^2^) was performed. Full acquisition parameters for all diffusion protocols are given in Supplementary Table 3.

### Quantitative multi-contrast acquisition

Quantitative multi-contrast mapping was performed by adding quantitative T_2_ and B_1_^+^ mapping to multi-shell diffusion and quantitative T_1_ mapping for one subject, all at the same 1.8 mm resolution. Quantitative T_2_ and B_1_^+^ mapping were performed at 1.8 mm by relaxometry on a range of TE’s (from 28.3 to 88.3 ms in steps of 20.0 ms), and by cosine model fitting of Exc-Sto-Rec = α°-2α°-α° for a range of multipliers α (from 30° to 110° in steps of 20°; Lutti et al., 2012), respectively. Although neither qT_2_ nor B_1_^+^ mapping benefits in terms of acquisition speed from MESMERISED’s echo-shifting or block reordering mechanisms, T_2_ relaxometry was performed at both ES = 1 and at ES = 3, to assess whether robust qT_2_ mapping can be performed under echo-shifting. Full acquisition parameters for qT_2_ and B_1_^+^ mapping are given in Supplementary Table 2.

### Image processing and analysis

Each MESMERISED dataset (all repetitions for all TM’s (qT_1_), TE’s (qT_2_), flip angles (B_1_^+^), or directions and b-values for diffusion) was individually denoised, separately for the SE volumes and STE volumes, except for the proof-of-principle signal comparison acquisitions (reported in Figure 2) and the MESMERISED – PGSE high-b diffusion comparison acquisitions (reported in Figure 9 and 10, as well as Supplementary Figures 2, 3 and 4). Denoising was performed before any other step following the approach by Veraart et al. (2016), using the mrdenoise tool in MRtrix 3.0 (Tournier et al., 2019) with a kernel size of 5 or 7. Subsequently, all data were corrected for movement, off-resonance and eddy-current artifacts using the Topup and Eddy tools of FSL v6.0.1 (Jenkinson et al., 2012). Given that the TE, EPI readout-train, and crusher/diffusion gradients are exactly matched between SE and STE within each acquisition, B_0_ field related distortion estimation can be performed on either SE or STE. Therefore, topup was performed on the highest signal SE volumes (i.e. shortest TE, flip angles 90° - 180° - 90°, and/or b_0_ acquisition). Since (particularly long time constant) eddy currents can differ between SE and STE, FSL Eddy was then performed separately on SE and STE volumes with this Topup input. Since data was acquired with shortest-TM volumes first, the first acquired SE and STE volumes are already in close spatial alignment (having been acquired ∼ 100 - 300 ms apart in time), which ensures good alignment of all SE and STE volumes. Brain masks for each subject and each MR contrast, needed for the preprocessing and analysis steps, were obtained using the BET tool of FSL v6.0.1. After correction, the following analyses were performed.

**Figure 2:**
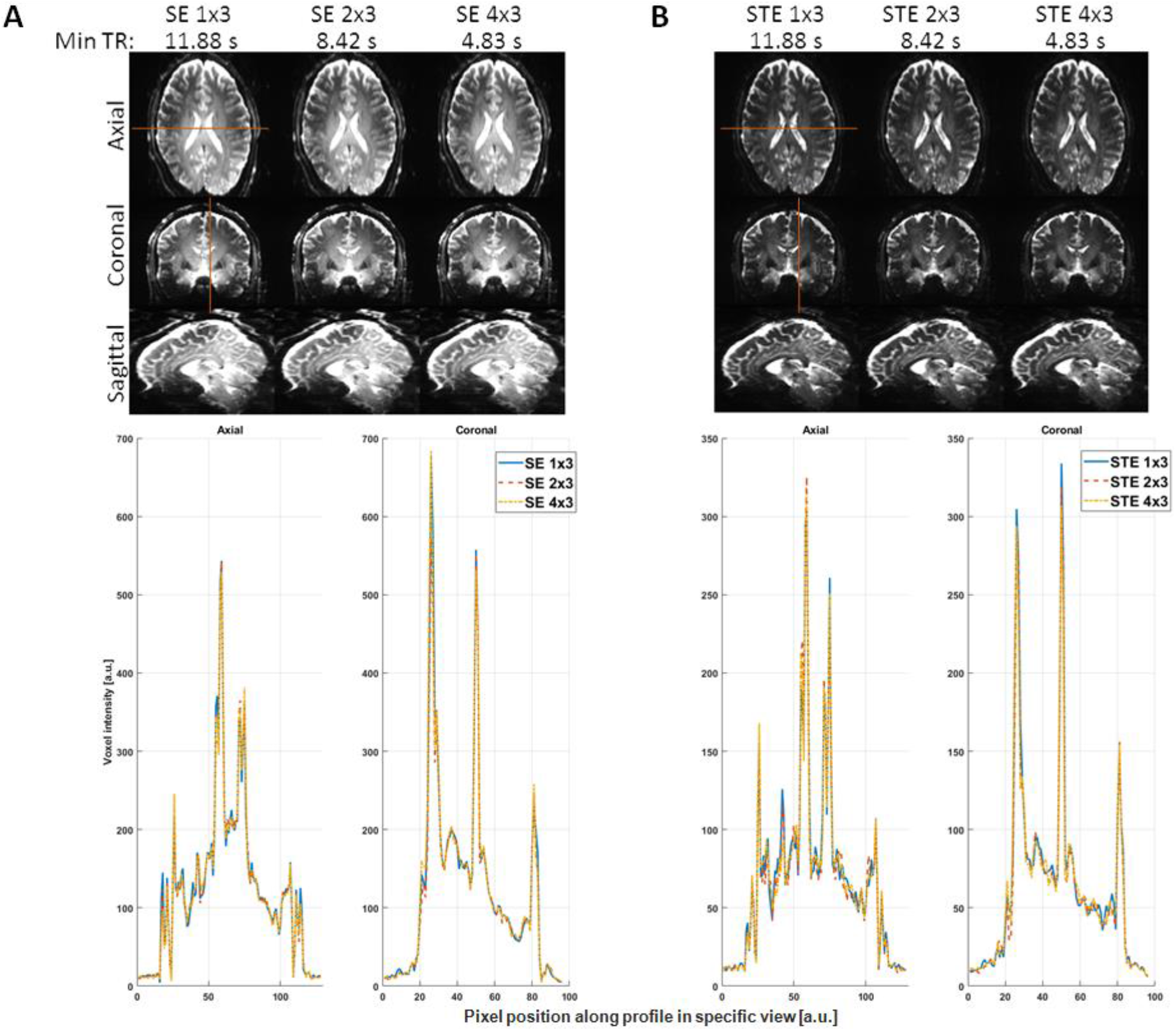
Comparison of MESMERISED images at 1.8 mm isotropic, acquired with MB = 3 at different echo-shift factors of ES = 1, ES = 2 and ES = 4 but otherwise identical parameters (TR = 11.88 s; TE = 47.5 ms; TM = 300 ms) for spin-echo (A, top) and stimulated echo (B, top) The minimal achievable TR (min TR) is indicated to illustrate increased acceleration capability with ES > 1. Bottom row: corresponding signal line plots for SE (A) and STE (B) for the three echo-shifts for the red line in the axial view (left line plot) and coronal view (right line plot).

### qT_1_ analysis

For qT_1_ analysis, two distinct relaxometry approaches were used which fit volumes at varying TM with the same TR and TE. In the first approach, only the STE volumes are used. The STE signal as a function of TR, TM, TE and flip angles can be written:

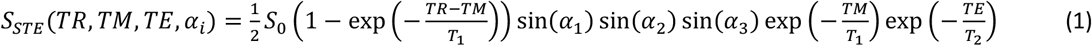

Where S_0_ contains all parameters not varying with TR, TM, TE and flip angles, such as proton density and Tx/Rx spatial sensitivity profiles, and α_1_, α_2_, and α_3_ are the flip angles (in radians) of excitation, storing, and recalling pulse, respectively. Absorbing all terms that do not vary with TR or TM into the *S*_*0*_ term, we get:

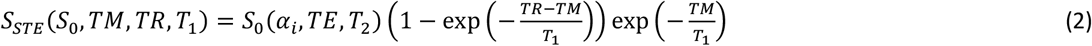

In the first approach, equation 2 was fitted to STE volumes at varying TM and maps for *S*_*0*_ and T_1_ were obtained. In this approach, the *S*_*0*_ absorbs effects of flip angle and T_2_ decay (as well as RF receive coil sensitivities and proton density), and the fitted curve is a product of exponentials.

In the second approach, both SE and STE volumes are used. The SE signal and STE/SE signal ratio as a function of TR, TM, TE and flip angles can be written:

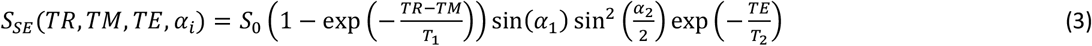

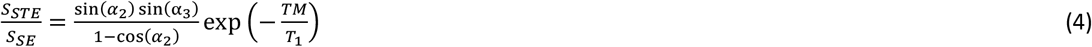

Absorbing all terms that do not vary with TR or TM for the ratio into an *S*_*0*_ ^*\**^ term, we get:

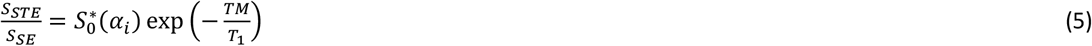

In the second approach equation 5 was fitted to STE/SE ratio volumes at varying TM and maps for *S*_0_ ^*\**^ and T_1_ were obtained. In this approach, the *S*_0_^*\**^ absorbs effects of flip angle, and the fitted curve is a simple exponential. Therefore, the second approach uses the available SE images in MESMERISED to establish a potentially lower complexity fit (simple exponential vs. product of exponentials in the first approach). Approach 1 and 2 were both evaluated for the estimation of qT_1_ maps and compared for the precision of their estimates by evaluating Fisher Information Matrix (FIM) local uncertainty (Harms et al., 2019). In addition, the uncertainty of qT_1_ estimates was evaluated for an increasing number (1, 3, 5) of repetitions per TM volume.

### Diffusion analysis

The diffusion-weighting in MESMERISED is the same as for the underlying diffusion-weighted STEAM sequence: whereas the SE signal is diffusion-weighted by gradients before and after the storing pulse, separated by the diffusion time Δ_SE_ (S_dSE_), the STE signal is weighted by the diffusion gradient before the storing pulse and the gradient after the recalling pulse, separated by the diffusion time Δ_STE_ (S_dSTE_; see Figure 1). The latter diffusion time is equal to Δ_SE_ + TM. The corresponding diffusion-weighted signal equations for both echoes are defined as:

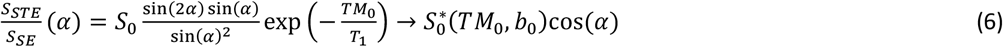

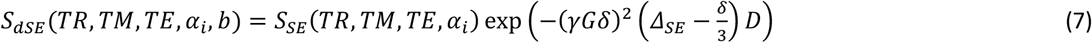

Where, S_SE_ and S_STE_ are the SE and STE signals defined in equations 1 and 3, respectively, γ is the gyromagnetic ratio (in Hz/mT), *δ* is the diffusion gradient duration (in s), *G* is the diffusion gradient amplitude (equal for the gradients-pair in the storing pulse and the gradient after the recalling pulse, in mT/m) and *D* is the (scalar) diffusivity (in m^2^/s). For diffusion MRI modeling the exponential diffusion terms in eq 7 and 8 were replaced with the desired diffusion model, with the corresponding b value and diffusion times. For the 4-shell (max b = 7000 s/mm^2^) single-TM acquisitions, the diffusion data from the STE signal was analyzed by DTI on the 1^st^ shell, Diffusion Tensor Kurtosis (Jensen et al., 2005) on the 1^st^ and 2^nd^ shell, Ball&Stick models (Behrens et al., 2007; Behrens et al., 2003) with one (B&S_r1_), two (B&S_r2_) and three (B&S_r3_) sticks on all shells, NODDI model (Zhang et al., 2012) on the 1^st^ and 2^nd^ shell, CHARMED model (Assaf and Basser, 2005) with one (CHARMED_r1_) and two (CHARMED_r2_) cylinder compartments on all shells. For the SE volumes from the same acquisitions, the highest b-value shell was analyzed with DTI. For the 2-shell multi-TM acquisition, both DTI (1^st^ shell) and DKI (1^st^ and 2^nd^ shell) analysis were applied to the STE signal.

For the MESMERISED-PGSE b = 7000 s/mm^2^ diffusion imaging comparison the SE and STE volumes of each MESMERISED run (ES = 2, 3, 4, and 6) and the PGSE volumes were each separately preprocessed for distortions and movements as described above, without denoising. Subsequently, the Ball&Stick model with two sticks (B&S_r2_) was fitted to the b = 7000 s/mm^2^ MESMERISED STE and PGSE volumes (60 directions + 6 b_0_s) and the Diffusion Tensor model was fitted to the MESMERISED SE volumes, evaluating voxel-wise uncertainty as described below. For consistency and completeness, the Tensor model was fitted to all SE volume series, despite the fact that the b-values (752, 483, 350, and 223 s/mm^2^ for ES factor 2, 3, 4 and 6, respectively) of the higher ES factors is likely too low for robust fitting of the Tensor model. The fraction-of-sticks (FS) map and fractional anisotropy (FA) maps, and their uncertainty, were used for assessment of the fit quality of the B&S_r2_ and Tensor models, respectively. Voxel wise temporal signal-to-noise ratio (tSNR) maps were calculated from the b_0_ only acquisitions, discarding the first 2 and last 5 of 27 volumes, computing tSNR = mean/std over the remaining 20 volumes, and lightly smoothing the resulting map with a Gaussian smoothing kernel with standard deviation of 0.5 voxel. Voxel wise tSNR efficiency (tSNR-eff) was computed as tSNR/sqrt(T_vol-acq_) and tSNR per unit time as tSNR/T_vol-acq_, where T_vol-acq_ is the single volume acquisition time, which is TR/2 for STE and SE in MESMERISED and equal to TR for PGSE. White matter (WM) summary statistics of tSNR-eff, tSNR per unit time and diffusion modeling uncertainty were computed over the same deep WM mask for all acquisitions, obtained from a thresholded FS map processed with a 3D image erosion and median filtering step to avoid effects of tissue boundaries or volume misalignment. The mean and standard deviation over the WM mask was computed for tSNR-eff, tSNR per unit time and diffusion modeling uncertainty, excluding voxel with FS or FA values lower than 0.15, since uncertainty estimation can be less reliable close to hard parameter value limits (Harms et al., 2019).

### B_1_+ and qT_2_ analysis

For B_1_^+^ mapping, the approach used by (Lutti et al., 2012) was used on the acquired data at multiple flip angles. It consisted in taking the ratio between STE and SE signals, where the flip angle ratio between the three STEAM pulses is *α*°-2*α*°-*α*° (e.g. for *α* = 90°: 90°-180°-90° degrees for excitation, storing and recalling pulses respectively). The STE/SE ratio is proportional to the cosine of the excitation flip angle:

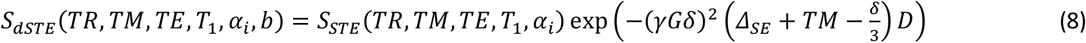

Therefore, the relative B_1_^+^ map can be estimated from a sufficient number of ratio volumes with varying *α*. For qT_2_ mapping only the SE signal was used for both acquisitions (ES = 1 and 3) separately, fitting a simple exponential decay of TE’s to estimate a qT_2_ map.

### Model optimization and uncertainty

All modeling analyses and parameter uncertainty quantifications above were performed in the Microstructure Diffusion Toolbox (MDT; https://mdt-toolbox.readthedocs.io/en/latest_release/) with Powell optimizer at patience 10, and using a cascaded optimization approach (Harms et al., 2017). For diffusion modeling, the cascade consisted of: S_0_ estimation – B&S_r1_ estimation – desired model; for qT_1_, qT_2_ and B_1_^+^ analysis, the cascade consisted of: S_0_ estimation – desired model. The precision or uncertainty of estimates was evaluated by computing standard deviations derived from the Fisher Information Matrix (FIM), tightly related to the Cramer Rao Lower Bound (CRLB). Whereas the CRLB amounts to the FIM evaluated around a ground truth value, the FIM can be evaluated around a maximum likelihood point estimate and this allows for estimating the standard deviation, or uncertainty, of the point estimate, both for linear (Andersson, 2008) and non-linear models (Alexander, 2008; Harms et al., 2019). FIM uncertainty was calculated as described in Harms et al. (2019) using numerical evaluation of the FIM, and analytic uncertainty propagation for the derived FA parameter for the Tensor diffusion model, as implemented in MDT.

## Results

Figure 2 shows a comparison between images obtained using MESMERISED without ES (ESxMB = 1×3, equivalent to MB-STEAM, cf. Figure 1A), ESxMB = 2×3 and 4×3 for the SE and STE images. Qualitatively, Figure 2 shows that images retain the same contrast and characteristics when dead time is filled up through echo-shifting. At the same time, the minimum TR (minTR) illustrates that MESMERISED allows these images to be acquired at higher acceleration in a shorter time, going from a 11.88 s minTR with ESxMB = 1×3, to 4.83 s minTR with ESxMB = 4×3. Here we should note that, if the shorter TR is actually used, the contrast and SNR might change compared to the long TR images shown here due to magnetization and steady-state effects. The minTR also illustrates that although the additional acceleration compared to multiband STEAM is significant (e.g. 2.46x for ESxMB = 4×3 compared to ESxMB = 1×3), and multiplicative with MB factor (e.g. 7.38x for ESxMB = 4×3 compared to ESxMB = 1×1), the temporal acceleration factor is generally not linear with ES, due to the time added by the STE_i_ (i> 1) blocks (c.f. Figure 1B). The bottom row of Figure 2 shows a more direct comparison in signal line plots. MESMERISED accelerated signals (2×3 and 4×3) follow the MB STEAM signal (1×3) closely.

### qT_1_ mapping

Figure 3A shows MESMERISED T_1_ relaxometry data at 1.8 mm isotropic. The relaxometry on TM is performed at constant TE (28 ms) and TR (6.9 s) by increasing the ES factor proportionally to TM, for TM’s from 140 ms to 1680 ms (see Supplementary Table 1). This results in total acceleration factors (ESxMB) from 3 to 36, highlighting the super-acceleration capability of MESMERISED for T_1_ relaxometry. For MB-STEAM and STEAM (without MB) the TR for relaxometry would be limited by the largest TM and would equal 41.51 and 124.60 s, respectively, which means that relative temporal speed-ups of 6.02x and 18.06x are achieved compared to MB-STEAM and STEAM, respectively. Figure 3B shows the whole-brain isotropic resolution S_0_ and qT_1_ maps resulting from analysis of STE data (i.e. using Equation 2). The S_0_ map estimated in this way retains strong T_2_ weighting (along with weighting for proton density and Tx/Rx fields), which is apparent in the contrast between white matter (WM) and gray matter (GM) structures. The qT_1_ map also shows a high contrast level due to the quantitative differences in T_1_ between WM and GM structures. For comparison, Figure 4A shows S_0_ ^*^ and qT_1_ maps resulting from analysis of the STE/SE data from the same acquisition (i.e. using equation 5). The resulting qT_1_ map is qualitatively highly similar. However, the theoretical expectation (described above) that the S_0_ ^*^ map is more weighted for the transmit field (i.e. local flip angle) and less for T_2_ compared to the S_0_ map is borne out in a decreased WM/GM contrast and an increased central darkening strongly (anti-)correlated with the transmit field (described below).

**Figure 2:**
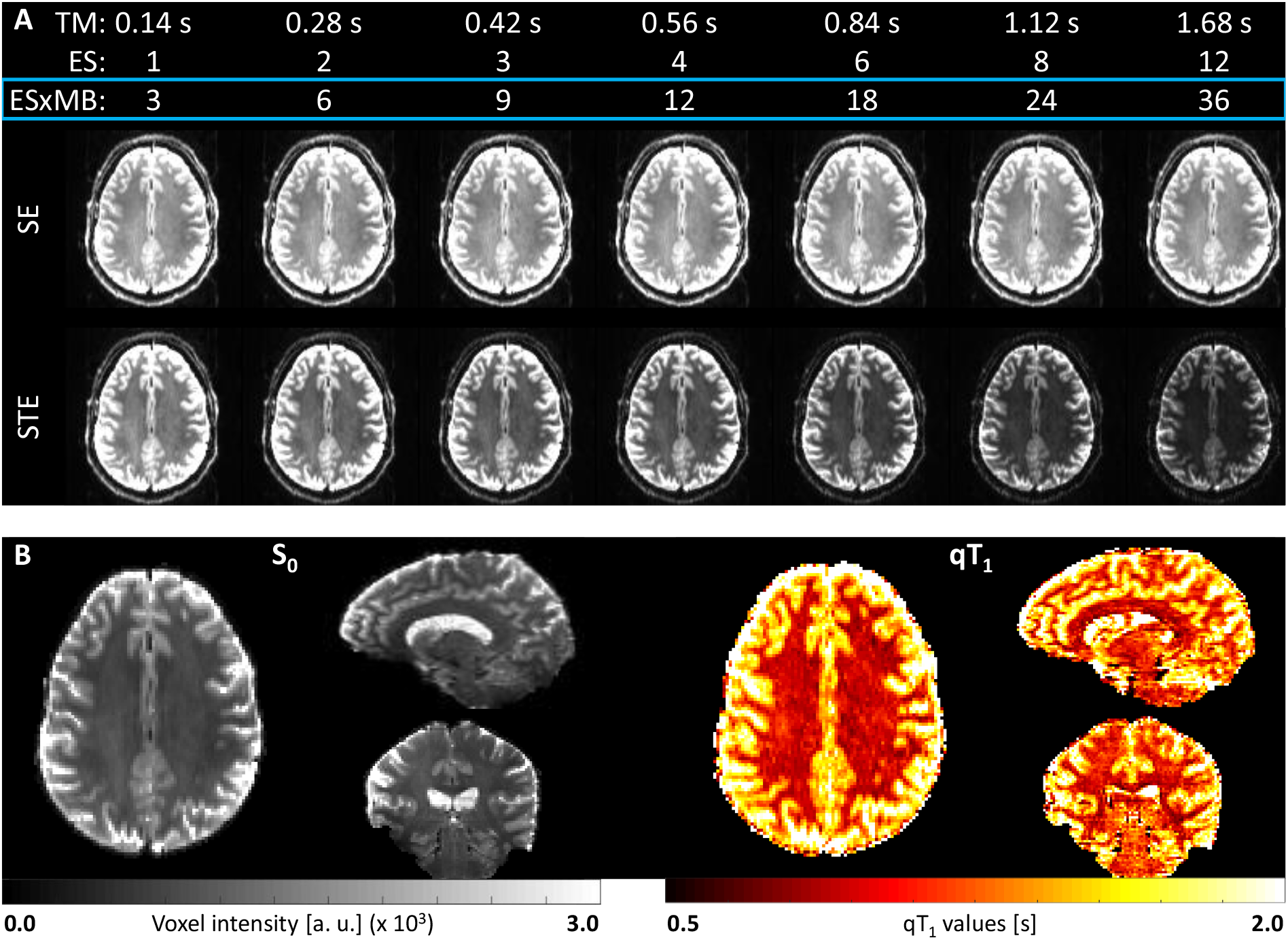
MESMERISED qT_1_ mapping and T_1_w relaxometry data at 1.8 mm isotropic. (A) Highly accelerated T_1_w data was acquired by increasing ES factor in tandem with TM to keep the TR constant and short TM, ES and total acceleration factor EsxMB are given. The acquired SE and STE signals are shown with the STE signal visibly attenuating with TM. (B) The corresponding S_0_ (left) and qT_1_ (right) maps estimated from the STE signal only (Equation 2).

**Figure 4:**
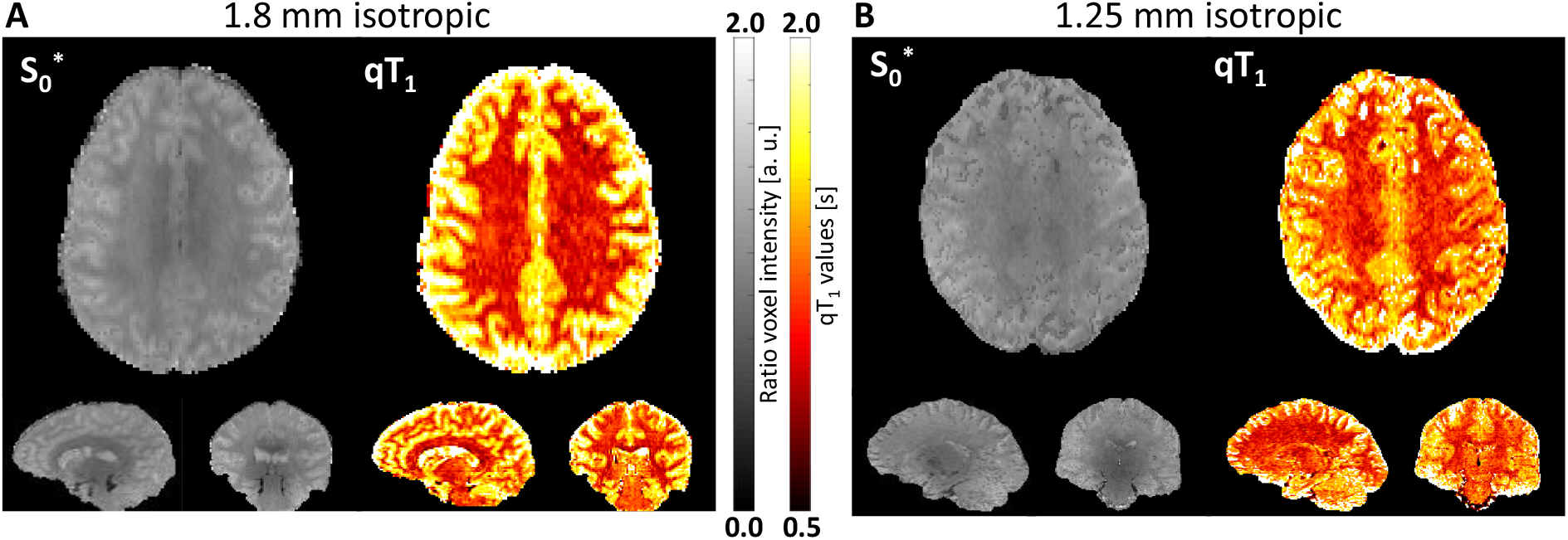
MESMERISED qT_1_ mapping at 1.8 mm (A) and 1.25 mm (B) isotropic using the STE/SE ratio T_1_-relaxometry model (Equation 5) on data acquired using 3 repetitions (TA: 2:07 and 4:01 min, respectively).

Figure 4B shows MESMERISED S_0_ ^*^ and qT_1_ maps at 1.25 mm isotropic, from data at constant TE (31.7 ms) and TR (8.8 s), resulting in total acceleration factors (ESxMB) from 3 to 54 over 8 different TM’s (Supplementary Table 1). For MB-STEAM and STEAM (without MB) the TR for relaxometry would be limited by the largest TM and would equal 79.79 and 239.40 s, respectively, which means that relative temporal speed-ups of 9.07x and 27.20x are achieved compared to MB-STEAM and STEAM, respectively.

A more detailed comparison of the uncertainty of qT_1_ maps for the more SNR challenged 1.25 mm isotropic data is shown in Figure 5. This compares a full duty cycle acquisition (cf. Figure 1C) with a maximum TM of 1080 ms (short TR, TM; top) with a half duty cycle acquisition were TR and all TMs are doubled (long TR, TM; bottom; maximum TM of 2160 ms). This shows that uncertainty in qT_1_ estimates is considerably lower for all long TR,TM acquisitions compared to short TR,TM acquisitions. Furthermore, an upward bias in qT_1_ estimates is visible for full duty cycle short TR,TM acquisitions. This is easily explainable by the TM range, implied by the block reordering property under full duty cycle (i.e. short TR,TM), that is insufficient (from 70ms to 1080ms) for the expected T1’s at 7 T (1 - 1.5 s). When TR and all TMs are doubled, and hence acquisition duty cycle is decreased to 50%, this range (140ms to 2160ms) is much more suitable and results in good estimates and uncertainty. Therefore, focusing on the bottom row, it is observed that uncertainty in qT_1_ estimates is considerably lower 1) with an increasing number of repetitions and 2) with STE/SE ratio analysis compared to STE-only analysis. Although 1) is expected given the averaging effect of repetitions, the benefit of STE/SE ratio analysis is more interesting, especially in comparison with the effect of repetitions. With the STE/SE ratio analysis the local maxima in uncertainty for the long TR, TM data is at a similar level with a single repetition (acquisition time 1:40) as for the STE-only analysis with 5 repetitions (acquisition time 6:22); the STE/SE ratio analysis with three repetitions (acquisition time 4:01) has lower uncertainty everywhere (Figure 5, bottom row). Additionally, the uncertainty for STE/SE ratio analysis is more homogenous over the brain. Therefore, the 1.25 mm isotropic whole-brain qT_1_ map in Figure 4 can be achieved in 2-4 minutes, and the 1.8 mm isotropic qT_1_ map in 1-2 minutes, depending on the desired precision level and STE/SE ratio analysis can provide higher precision, comparable to that achieved by averaging over multiple repetitions.

**Figure 5.**
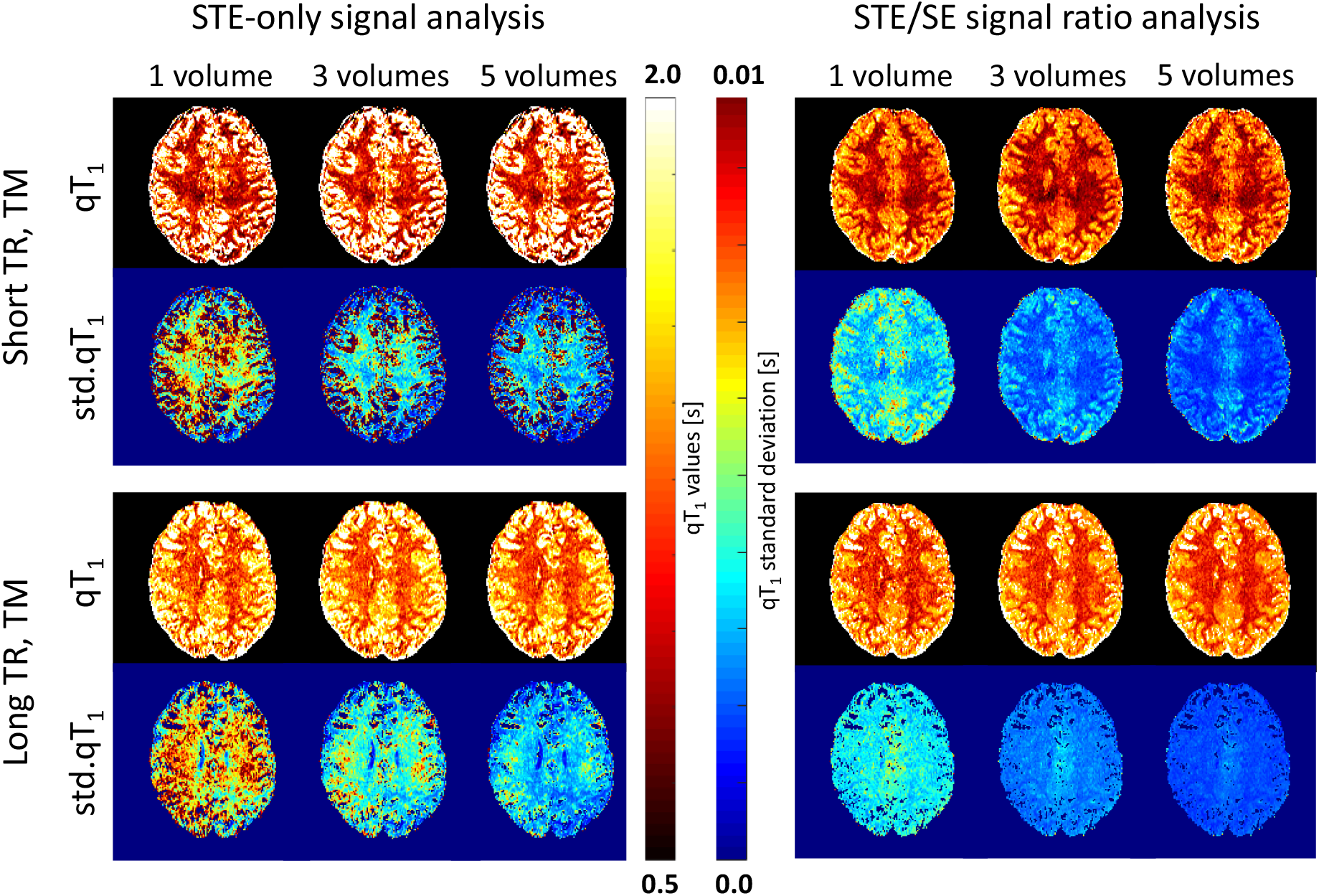
MESMERISED 1.25 mm isotropic qT_1_ mapping and its uncertainty compared between: short vs. long TR and TM (top vs. bottom), STE-only vs STE/SE ratio analysis (left vs. right) and number of repetitions (1, 3 and 5). Short TR is 4.4 s with a maximum TM of 1080 ms, in which case 1/3/5 repetitions have an acquisition time of 1:05 / 2:16 / 3:26 respectively; long TR is 8.8 s with a maximum TM of 2160 ms, in which case 1/3/5 repetitions have an acquisition time of 1:40 / 4:01 / 6:22 respectively. STE-only qT1 analysis is performed with Equation 2, whereas STE/SE ratio analysis is performed with Equation 5. Uncertainty or precision of the point-estimates is mapped as standard deviations derived from the Fisher Information Matrix.

### Diffusion MRI

Figure 6 illustrates the two-dimensional parameter space of diffusion times (vertical) and b-value (horizontal) with MESMERISED diffusion data at 1.8 mm isotropic (where the further 2 dimensions of diffusion direction on the sphere are left implicit). Here, different diffusion times were realized by varying TM and utilizing the maximum echo-shift factor allowed at each TM, individually minimizing TE and TR. Both TR’s are close to 3.4 s (see Supplementary Table 3), showing a capacity to explore high b-values, up to 7000 s/mm^2^, and different diffusion times, at high acceleration and data rate. As before, the relative temporal speedup achieved with MESMERISED will increase with ES factor and here speedups are achieved compared to MB-STEAM and STEAM of 1.47x and 4.40x for ES = 2, and 1.95x and 5.85x for ES = 3, respectively. In the following, we first focus on the possibilities afforded by multi-shell acquisitions at a single diffusion time and then demonstrate the possibilities of acquisitions at varying diffusion times.

**Figure 6:**
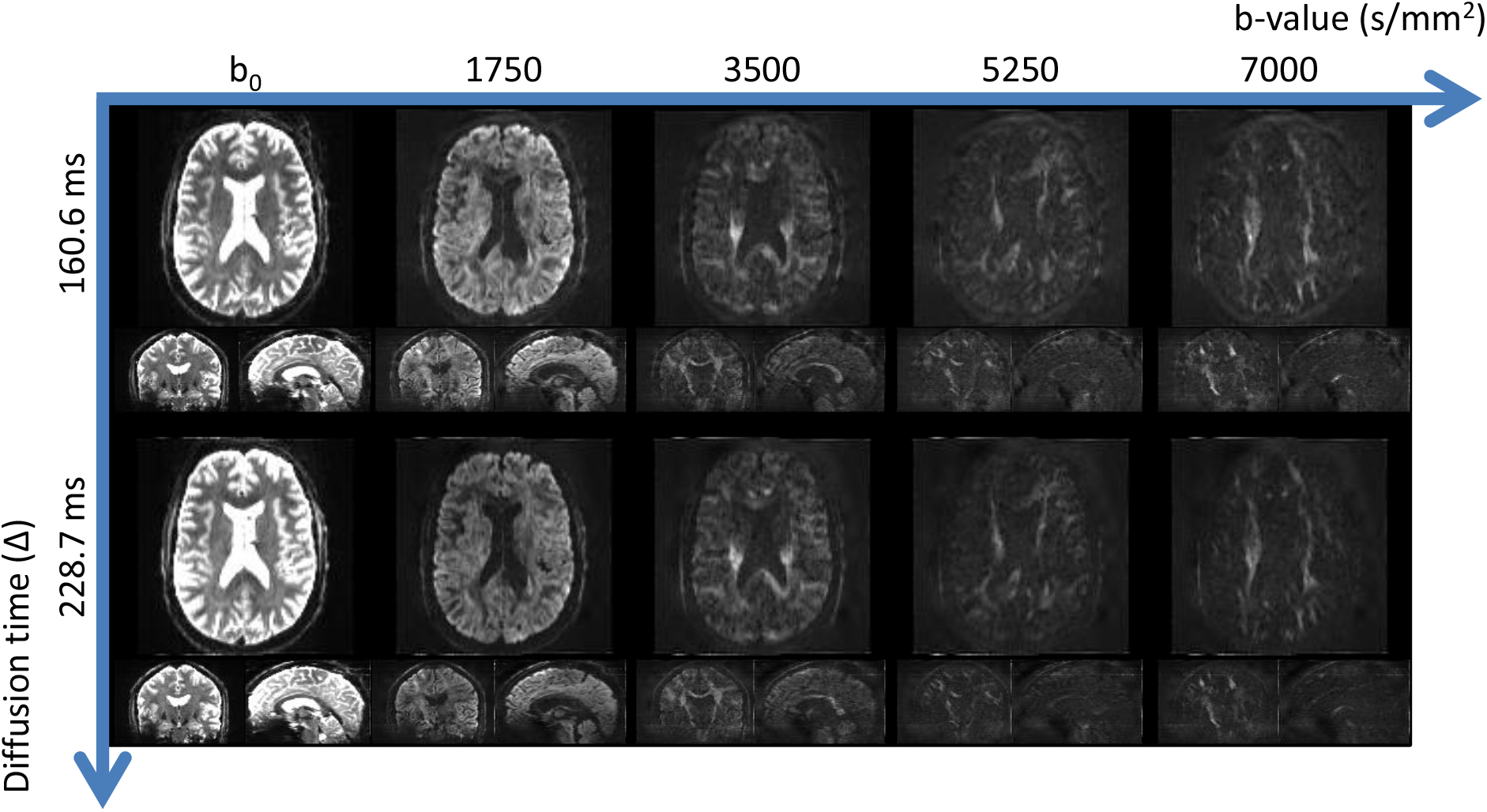
MESMERISED diffusion data at 1.8 mm isotropic spanning the space of diffusion time (vertical) and b-value (horizontal). Whole-brain STE images are shown for two different diffusion times, each corresponding to a distinct ES factor (2 for 160.6 ms and 3 for 228.7 ms) and individually optimized TE and TR, with different b-values from the same multi-shell acquisition. Simultaneously acquired SE images are not shown.

### Multi-shell diffusion MRI

Figure 7 shows example slices out of a 4-shell MESMERISED acquisition and analysis results for the simultaneously acquired STE images (at a diffusion time of 160.6 ms and b-values of 1750, 3500, 5250, and 7000 s/mm^2^) and SE images (at a diffusion time of 20.6 ms and b-values of 188, 376, 564, and 752 s/mm^2^). This makes explicit how the acquisition of both diffusion-weighted SE volumes and STE volumes provides 8 useful shells of diffusion data in a 4-shell acquisition, even when analyzed separately. Whereas the low-b SE volumes support analysis with Gaussian diffusion models such as the DTI model, the high-b STE volumes are suitable for non-Gaussian diffusion models, such as the DKI, NODDI and CHARMED models. Here, the Tensor model was applied in both the lowest b-value STE shell (Figure 7A) and the highest b-value SE shell (Figure 7B), showing some qualitative differences, likely related to the large difference in b-value (752 vs. 1750 s/mm^2^) at the limits of applicability of the Tensor model. This data was acquired at ES = 2 at a TR of 3.45 s and a TE of 49.8 ms, which tunes the diffusion time of the STE volumes to the shorter end of the achievable range and the b-values of the SE volumes to the higher end, which is accompanied with a higher TE than would be the case for higher echo-shift factors. When a longer diffusion time and higher echo-shift factor is chosen for the same STE volume maximum b-value (e.g. 7000 s/mm^2^ here), diffusion gradient duration d naturally decreases, along with the SE maximum b-value (see Supplementary Table 4).

**Figure 7:**
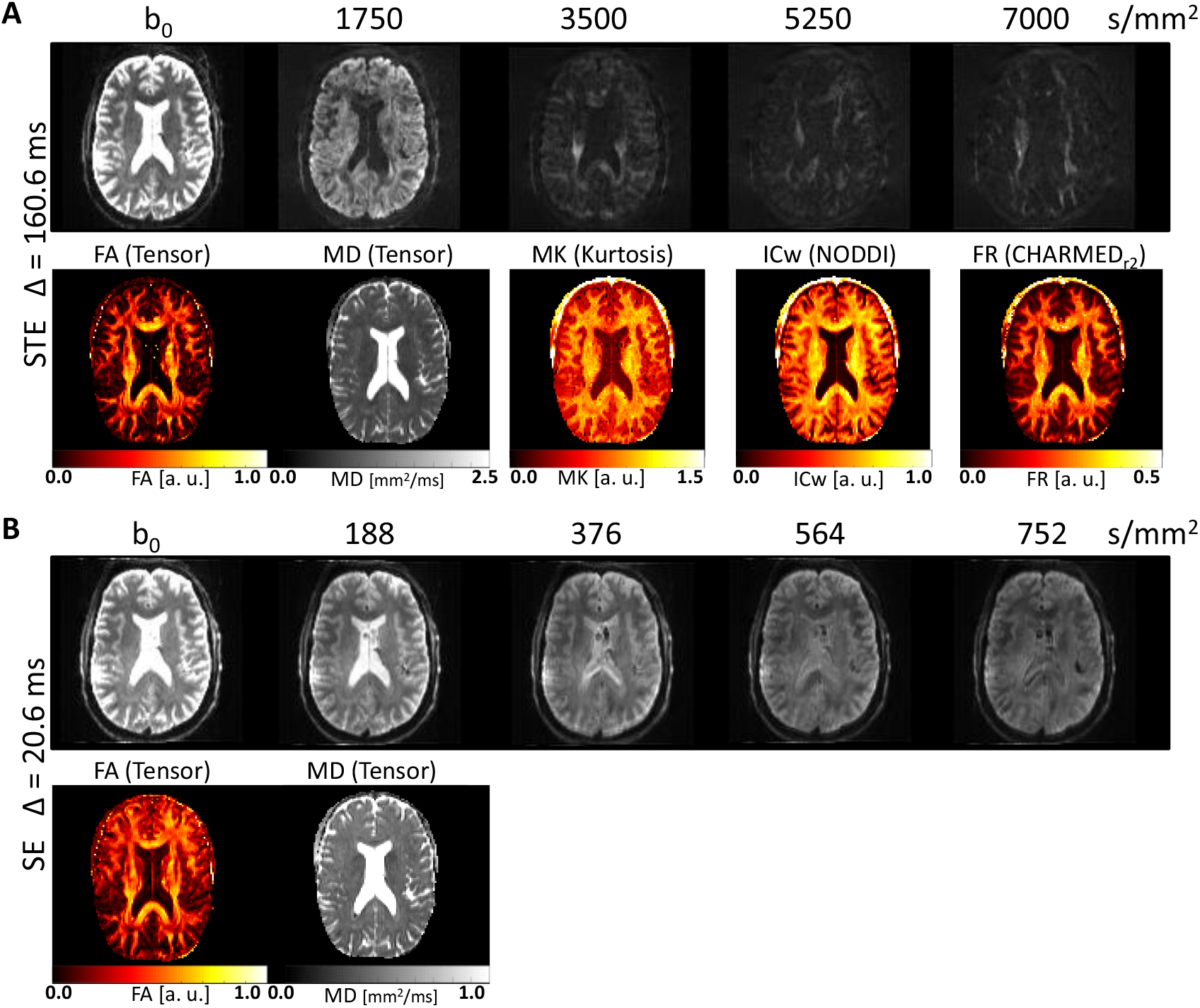
MESMERISED data and diffusion modeling results for simultaneously acquired STE volumes (at high b-values and diffusion time 160.6 ms) and SE volumes (at low b-values and diffusion time 20.6 ms) at 1.8 mm isotropic (ES = 2, TR = 3.45 s, TA = 11:08). (A) Example transverse slices through STE volumes at each b-value in the 4-shell (168 direction) acquisition (top) and their analysis with Tensor, Kurtosis, NODDI and CHARMED models (applied to suitable combinations of shells, bottom). (B) Example transverse slices through SE volumes at each b-value in the 4-shell (168 direction) acquisition (top) and their analysis with the Tensor model (bottom).

Figure 8 shows two 4-shell (max-b = 7000 s/mm^2^) acquisitions which realize full duty cycle at similar mixing and diffusion time at 2 mm isotropic (ES = 5, TM = 290 ms; top) and 1.8 mm (ES = 4, TM = 280 ms; bottom). The relative temporal speedup achieved compared to MB-STEAM is 2.95x for the 2 mm dataset and 3.43x for the 1.8 mm dataset. The 2 mm acquisition achieves a shorter TR (2.36 s vs. 3.37 s) and therefore enables a higher data rate of ∼ 1.2 s per volume (for SE and STE together). This generates the full 8-shell 370 volume dataset in an acquisition time of 7 min and 47 s (vs. 10 min and 53 s for 1.8 mm), showing that very short acquisition times are possible for extensive multi-shell data, particularly at resolutions of 2 mm and above, at which high b-value diffusion acquisitions are mostly performed.

**Figure 8:**
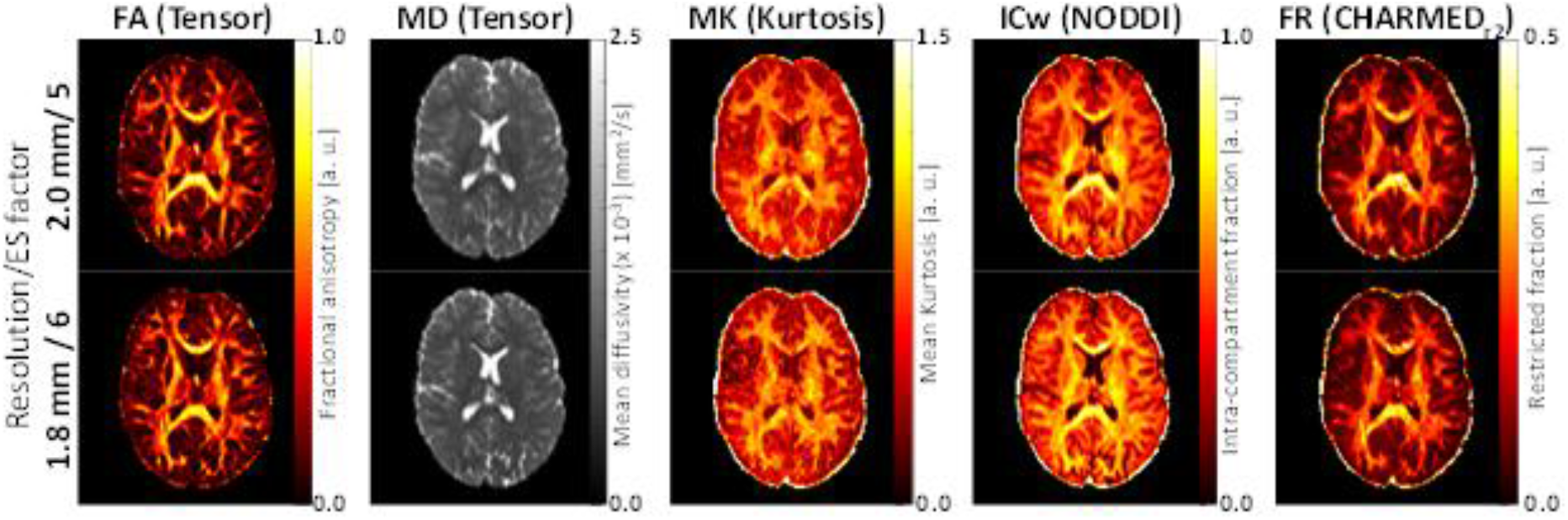
MESMERISED diffusion modeling results for 185 direction multi-shell MESMERISED diffusion acquisitions at different resolutions. Analysis results for Tensor, Kurtosis, NODDI and CHARMED models (applied to suitable combinations of shells) for 2.0mm isotropic (ES = 5, TR = 2.36 s, TA = 7:47) and 1.8 mm isotropic (ES = 6, TR = 3.37 s, TA = 10:53) in the same subject.

### Comparison with PGSE for 7 T high-b multi-shell diffusion MRI

We compared MESMERISED’s tSNR efficiency for high b-value 7 T multi-shell acquisitions with MB PGSE, focusing on the highest b-value shell (b = 7000 s/mm^2^) which is the most SNR limiting. Figure 9 shows results for MESMERISED for four different ES factors and at MB = 3 (Fig. 9A-D) and PGSE at MB = 2 (Fig. 9E), optimized for efficiency given SAR and gradient system limits. It can be seen that MESMERISED provides two useful volumes of multi-shell data (STE and SE) at two b-values in a shorter TR (∼3.4 s) than the TR for a single volume (4.21 s) for PGSE. In MESMERISED, tSNR is higher for SE than STE, as expected from the additional T_1_ decay in STE, and tSNR maps for MESMERISED STE show higher values in both deep and cortical gray matter than PGSE. However, since high b-value diffusion acquisitions are generally aimed at white matter, we quantitatively compared tSNR efficiency (tSNR-eff) in white matter. White matter tSNR-eff is markedly higher for both STE and SE volumes of all MESMERISED acquisitions than for PGSE (Fig. 9F, Table 1). MESMERISED STE tSNR-eff is at a stable level for all ES factors (2,3,4,6) while MESMERISED SE tSNR-eff increases with ES and decreasing TE. MESMERISED’s main efficiency advantage is through the acquisition of both STE and SE volumes in a shorter TR, providing a far shorter acquisition time per volume (1.7 s versus 4.21 s for PGSE). Thus, although MESMERISED STE white matter tSNR, without regard of acquisition time, is 8.4% higher than for PGSE (e.g. for ES = 3: 13.89 vs. 12.81), tSNR per unit time is 171% higher for MESMERISED (8.24 s^-1^ vs 3.04 s^-1^), and tSNR efficiency is 71% higher for MESMERISED (10.70 s^-1/2^ vs 6.24 s^-1/2^; Table 1).

**Figure 9:**
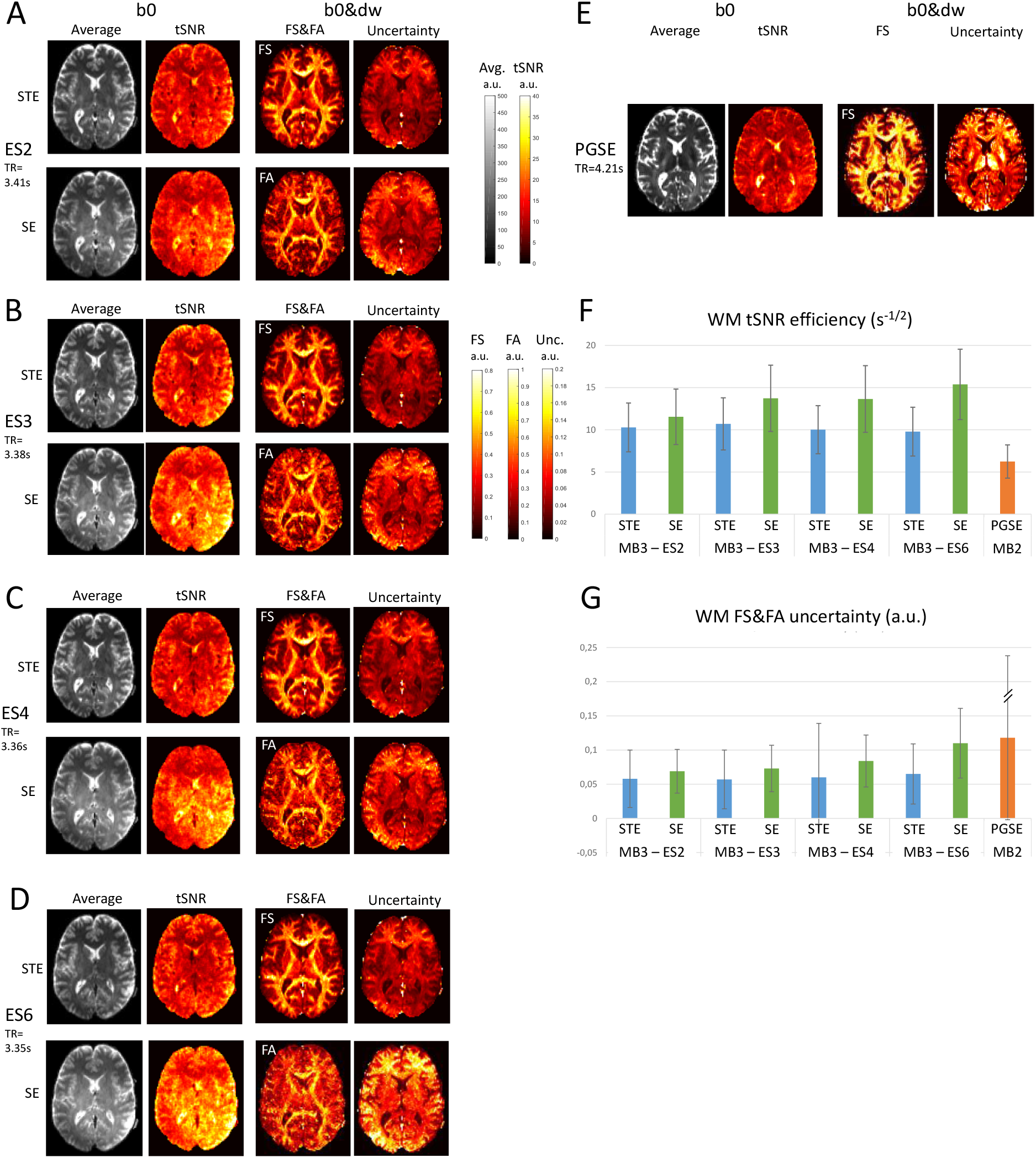
A comparison of MESMERISED with PGSE for high b-value multi-shell diffusion MRI at 7 T. (A-D) For MESMERISED with each of ESxMB = 2×3 (ES2 in A), ESxMB = 3×3 (ES3 in B), ESxMB = 4×3 (ES4 in C), ESxMB = 6×3 (ES6 in D): a transverse slice of whole brain 1.8mm isotropic results for MESMERISED with both STE (top) and SE (bottom) results. The left two panels show results derived from the b0 only acquisition with the average b0 volume and tSNR. The right two panels show results derived from the 60dw + 6b0 acquisition with FS (for STE) and FA (for SE) and their uncertainty. (E) PGSE results with MB = 2 as in A-D for its single spin-echo. (F) Mean tSNR efficiency over white matter only for all MESMERISED (MSMD) acquisitions (both STE and SE volumes) and PGSE. (G) Mean uncertainty over white matter only, for all MESMERISED (MSMD) acquisitions (both STE and SE volumes) and PGSE. Error bars: standard deviation. //: uncertainty error bar for PGSE downscaled 30x. All map scaling equal, and according to the color scale bars.

**Table 1.**
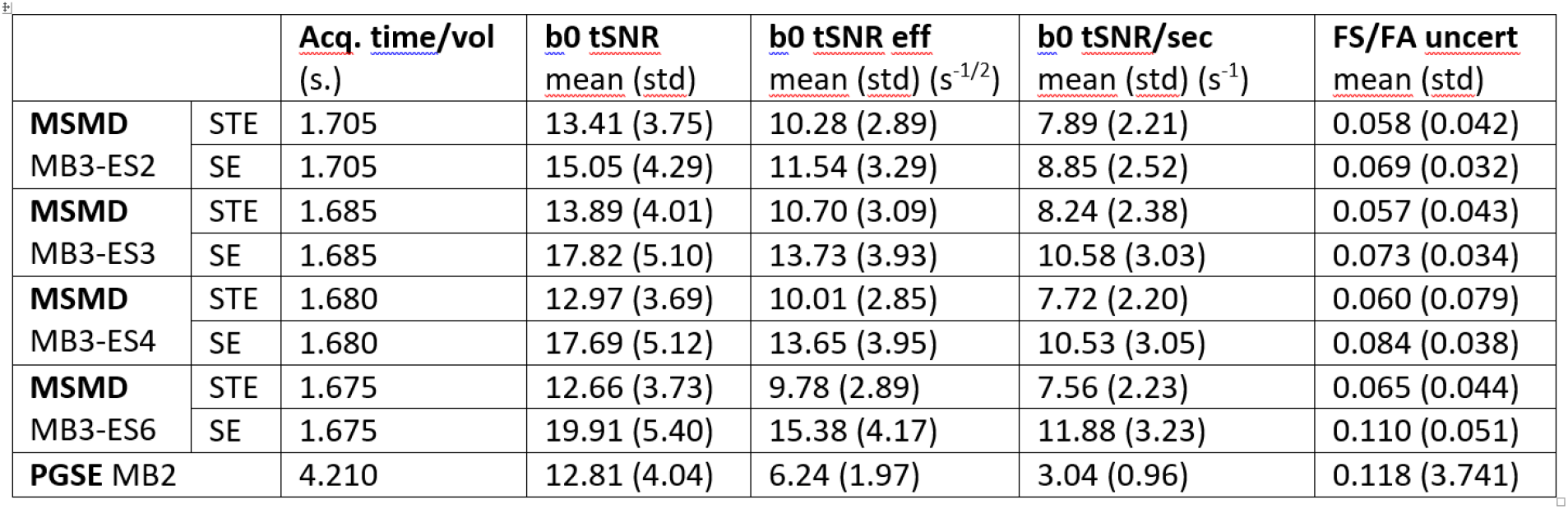
**Quantitative results for the comparison of MESMERISED with PGSE for b = 7000 s/mm2 diffusion imaging. Results for MESMERISED with MB = 3 and ES = 2, 3, 4, and 6, for each of the STE (b = 7000s/mm2) and corresponding SE volumes and for PGSE (b = 7000s/mm2) with MB = 2. Acq. time/vol: acquisition time per volume, TR/2 for MESMERISED STE and SE, TR for PGSE, as the basis for reported SNR efficiency and SNR per unit time calculations. b_0_ tSNR: temporal SNR calculated from b_0_ volumes, mean and standard deviation over white matter. b_0_ tSNReff: temporal SNR efficiency (in units of s^-1/2^) calculated from b_0_ volumes, mean and standard deviation over white matter. b_0_ tSNR/sec: temporal SNR per unit time (in units of s^-1^) calculated from b_0_ volumes, mean and standard deviation over white matter. FS/FA uncert: the uncertainty of fraction-of-sticks (FS for MESMERISED STE and PGSE volumes) and DTI fractional anisotropy (FA for MESMERISED SE volumes), mean and standard deviation over white matter.**

The standard deviations of 30 - 40% of the mean indicate that there is quite some variation of tSNR over white matter in all cases. A comparison of tSNR with a B_1_^+^ field map (Figure 10A, B) shows that B_1_^+^ inhomogeneity is likely a factor in this: the spatial tSNR variation follows similar low frequency patterns in STE, SE and PGSE, which co-vary with B_1_^+^ inhomogeneity. For instance, the region of low flip angle deviation at the green arrow on the subject’s left side (Figure 10A) is associated with high tSNR, whereas a region of high flip angle deviation, at a similar distance to the RF receiver coils (red arrow) shows low tSNR. With about 20% overflipping in the center of the brain, underflipping in regions of low B_1_^+^ reaches close to 50% (particular in right inferotemporal and inferior cerebellar regions; Supplementary Figure 2A,B), which are also the regions of lowest tSNR, up to 50% lower than highest observed values in each of MESMERISED STE, SE and PGSE (Supplementary Figure 2D, 5D and D6, respectively).

**Figure 10:**
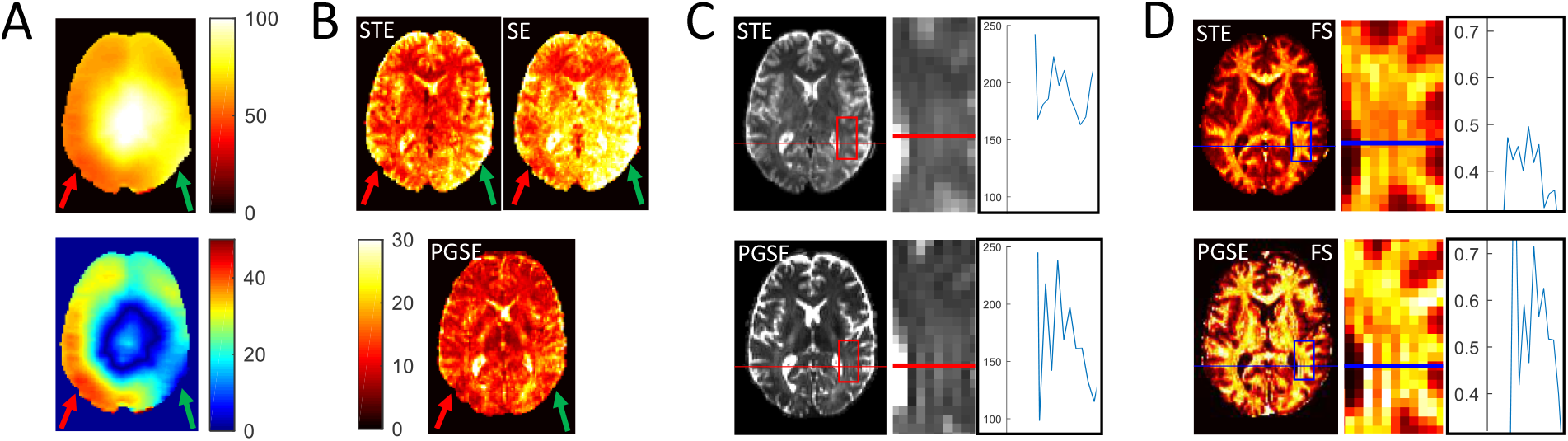
7 T B_1_^+^ field inhomogeneity and Gibbs ringing in high-b diffusion imaging with MESMERISED and PGSE. (A) A transverse slice through an absolute B_1_+ map (in units of degrees; top) and the deviation of the local flip angle from the nominal 90 degrees (bottom) with indication of regions of low deviation (green arrow on the subject’s left side; radiological display convention) and of high deviation (red arrow for the subject’s right side). (B) tSNR for MESMERISED with ESxMB = 3×3 for both STE and SE b0 volumes and PGSE. Regions of low and high flip angle deviation are marked as in A. (C) The mean b_0_ image for MESMERISED (ESxMB = 3×3) STE volumes and PGSE. Magnified portion at the edge of the lateral ventricle (middle; position indicated by the red box) and line-plot of single voxel intensities (right; position indicated by the red line). (D) The Fraction of Sticks (FS) map for MESMERISED (ESxMB = 3×3) STE volumes and PGSE. Magnified portion at the edge of the lateral ventricle (middle; position indicated by the blue box) and line-plot of single voxel intensities at the edge (right; position indicated by the blue line).

Beyond tSNR and tSNR efficiency, we assessed the effectiveness of MESMERISED and PGSE to support fiber orientation and microstructure modeling (Figure 9A-E, right columns, and used the uncertainty of FS and FA as a measure of modeling precision supported by the data. Fraction-of-stick (FS) maps for the high-b MESMERSED STE volumes have excellent WM-GM contrast for all ES factors. In clear distinction, FR for PGSE lacks WM-GM contrast, particularly in deep GM nuclei, and appears to have an overall upward bias. Fractional anisotropy (FA) maps for low-b MESMERISED SE volumes also show good delineation of WM and GM for ES = 2, 3 and 4. Overall, FS and FA uncertainty and tSNR appear negatively associated and driven by B_1_^+^ inhomogeneity within each volume, a pattern which is evident over the whole brain (Supplementary Figures 2-4). FS uncertainty for STE appears lowest of all maps and relatively constant over all ES factors. FA uncertainty for SE increases with ES factor, despite increasing tSNR, likely reflecting the decreasing b-values with ES (cf. supplementary table 4). In particular, the high ES = 6 FA uncertainty values likely reflect the low b = 223 s/mm2 associated with these SE volumes, which is very low for DTI modeling, although low uncertainty values are still observed in regions of high tSNR. Compared over white matter, FS uncertainty over white matter is markedly lower for MESMERISED STE volumes than for PGSE, and white matter FA uncertainty is relatively low for MESMERISED SE volumes for ES = 2, 3, and 4 (Figure 9G). A significant factor beyond tSNR in the upward bias and higher uncertainty of FS for PGSE could be Gibbs ringing. Indeed, line plots of PGSE signal at the border of the lateral ventricle show considerable ringing which is much decreased in MESMERISED under the same image reconstruction (Figure 10C), corresponding to water signal that is roughly twice as high in PGSE with roughly similar WM signal (Supplementary Figure 2C vs 4C). The PGSE signal ringing clearly translates to strong ringing the PGSE FS map, whereas ringing in the MESMERISED FS map is strongly decreased in comparison (Figure 10D).

### Varying diffusion time

Analogous to keeping TR constant in MESMERISED T_1_ relaxometry (by increasing ES factor proportionally to TM), different diffusion times for the STE volumes at the same TR can be realized by increasing ES factor with increasing diffusion time. When TE, b-value and diffusion gradient duration are also kept constant, this opens the possibility to explore the behavior of diffusion phenomena over different diffusion times. Figure 10 illustrates this with 2-shell diffusion data acquired at four diffusion times with otherwise matched parameters (i.e. TR, TE, δ and b-value are equal throughout, see Supplementary Table 3). Each 1.8 mm isotropic dataset had 24 directions at b = 1000 s/mm^2^, 48 directions at b = 2500 s/mm^2^ and 8 b_0_’s, for a total of 70 STE volumes (and 70 SE volumes, not shown here, with two-shell b-values for ES factor 1, 2, 3 and 4, respectively: [52, 130], [68 and 171], [99, 247], and [180, 450] s/mm^2^). Because of the same-TR requirement the relative temporal speedup for ES = 4 applies to the entire dataset and the speedup achieved compared to MB-STEAM is 2.38x, which means the < 17-minute acquisition time for MESMERISED would increase to over 40 minutes with MB-STEAM. The multi-shell data is sufficient for whole-brain DTI analysis (on the b = 1000s/mm^2^ shell, Figure 10B) and DKI analysis (on both shells) at each diffusion time (Figure 10C), and to assess how diffusion measures such as mean diffusivity (MD) and mean kurtosis (MK) change with diffusion time (Figure 10D).

### Quantitative multi-contrast mapping

STEAM’s ability to perform qT_2_ and B_1_^+^ mapping in addition to qT_1_ and diffusion imaging creates an opportunity for fast multi-contrast mapping. Therefore, we explored MESMERISED’s capacity for fast mapping of qT_1_, qT_2_, B_1_+ and high-b diffusion with the same pulse sequence, given that qT_1_ and diffusion are greatly accelerated. Here it is important to note that qT_2_ and B_1_^+^ mapping, per se, do not benefit from the acceleration by echo-shifting in MESMERISED. However, both these STEAM mapping methods do not require long TMs and are intrinsically already fast (and full acquisition duty cycle) techniques. Figure 12 shows whole brain quantitative multi-contrast mapping at 1.8 mm isotropic, achieving qT_1_, qT_2_, B_1_^+^ mapping as well as extensive diffusion mapping in about 17 minutes. Crucially, the use of the same sequence and readout EPI provides for intrinsically aligned maps. Supplementary Figure 5 shows how qT_2_ maps are derived from relaxometry on TE (both with and without echo-shifting), showing that qT_2_ mapping with the SE volumes can be performed independent of echo-shifting and the TM of the STE. Supplementary Figure 6 shows how MESMERISED B_1_^+^ mapping is derived from both the SE and STE volumes for different α_TX_ multipliers in an α_TX_ - 2α_TX_ - α_TX_ flip angle scheme for Exc – Sto – Rec pulses. The result in Figure 12 is for 3 repetitions of all volumes in the qT_1_, qT_2_ and B_1_^+^ mapping as well as a total of 8 b-shells over 370 volumes for the diffusion. This delivers high precision (low uncertainty, cf. Figure 5) in qT_1_, qT_2_ and B_1_^+^ mapping, each in roughly 2 minutes, and diffusion mapping in 11 minutes. Alternatively, acquiring a single repetition for the qT_1_, qT_2_ and B^+^ maps, each in roughly 1 minute, and acquiring 200 volumes for the diffusion dataset (cf. Supplementary Tables 1-3), would lower the total multi-contrast acquisition time to under 10 minutes (i.e. qT_1_ mapping in 1:18, qT_2_ mapping 0:56, B_1_^+^ mapping 1:05 and 200 volume multi-shell diffusion in 6:15).

**Figure 11:**
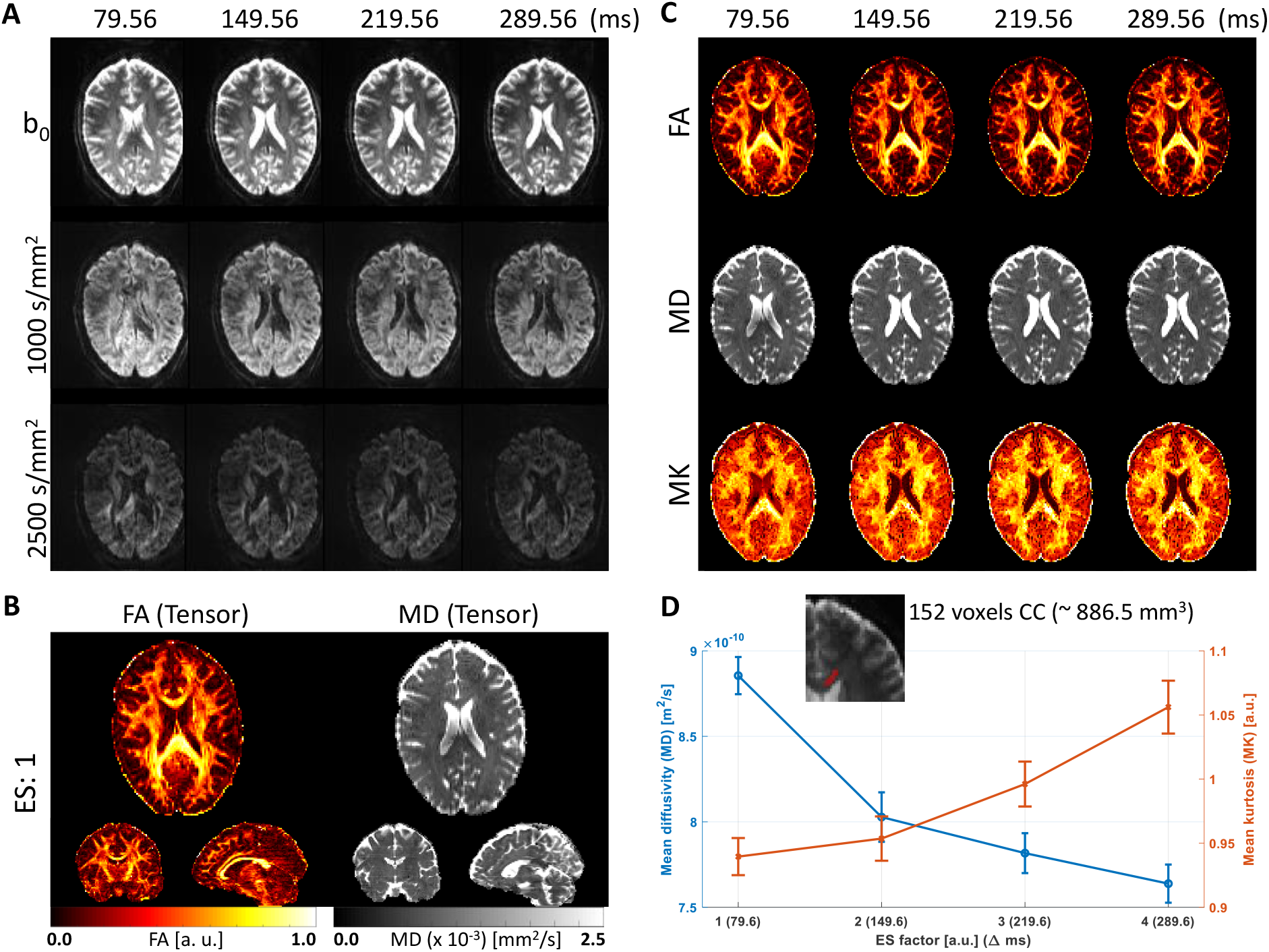
MESMERISED data and diffusion modeling results for STE volumes over several diffusion times at two b-values with matched TE and TR (TR = 3.50 s, TE = 45 ms) at 1.8 mm isotropic. (A) Example transverse slices through STE volumes at the two b-values and four diffusion times. (B) Whole-brain Tensor modeling resulting in FA and MD maps for the ES = 1 data (b = 1000 s/mm^2^ shell only). (C) Tensor and Kurtosis modeling results (applied to suitable combinations of shells) for the different diffusion times. (D) Mean diffusivity (MD, blue) and Mean Kurtosis (MK, red) with their corresponding standard error plotted over ES factors/diffusion times for an ROI (red in the inset, with a total of 152 voxels along 10 slices) in the genu of the corpus callosum.

**Figure 12:**
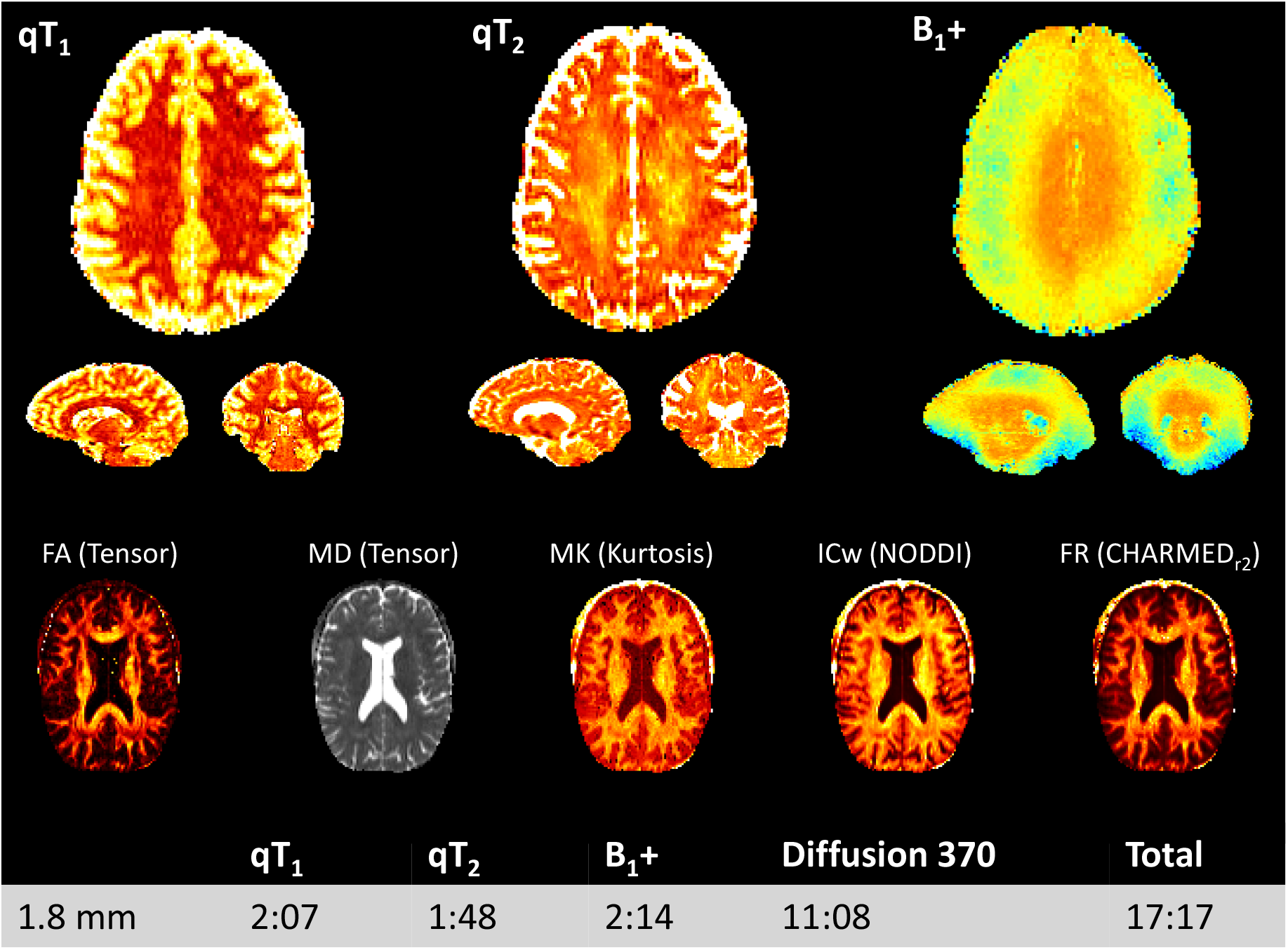
Whole brain quantitative multi-contrast mapping with MESMERISED at 1.8 mm isotropic in 17 minutes 17 seconds. This high precision protocol consists of 3 repetitions of all volumes in the qT_1_, qT_2_ and B_1_^+^ as well as 185 high-b multi-shell STE volumes and 185 low-b multi-shell SE diffusion weighted volumes for a total of 8 b-shells over 370 volumes.

## Discussion

MESMERISED achieves super-accelerated 7 T STEAM imaging by combining echo-shifting and multiband/simultaneous multi-slice acceleration, leading to very high multiplicative acceleration factors and time efficiency for qT_1_ and diffusion imaging. Furthermore, qT_2_ and B_1_^+^ mapping can be performed with the same pulse sequence. MESMERISED inherits from STEAM the capacity to simultaneous generate both SE images and STE images and the possibility of performing qT_1_, qT_2_, B_1_^+^ mapping as well as diffusion-weighted imaging. It also inherits the property of 50% lower SNR and 40% lower SAR compared to a similar spin-echo sequence, and 66% lower SAR compared to an inversion-prepared spin-echo sequence. In essence, MESMERISED capitalizes on the low SAR of the STEAM sequence to considerably increase its SNR per unit time through the multiplicative ESxMB acceleration. This enables fast qT_1_, diffusion and quantitative multi-contrast mapping over the whole-brain, which we will discuss in turn.

### qT_1_ mapping

MESMERISED T_1_ relaxometry can be performed at a low constant TR through the block reordering property (Figure 1): increasing the ES factor proportionally to TM results in increasing total acceleration factors (e.g. from 3 to 36 in 1.8 mm data and from 3 to 54 in 1.25 mm data in Figure 4), highlighting the super-acceleration capability of MESMERISED for qT_1_ imaging. Although the temporal speed-up of the multiplicative ESxMB acceleration is not linear with ES, the speed-up compared to multiband only accelerated STEAM can be very large (e.g. 6.02x faster than MB = 3 for 1.8 mm and 9.07x faster than MB = 3 for 1.25 mm). In addition, MESMERISED simultaneously provides spin-echo images in the same acquisition time which, we show here, can be used to either improve the precision of the qT_1_ maps in the same acquisition time or achieve even shorter acquisition times for the same precision level. For instance, uncertainty in 1.25 mm isotropic qT_1_ maps at a single repetition (acquisition time 1:40) with STE/SE ratio analysis are at similar levels as that of three repetitions (acquisition time 4:01) or even 5 repetitions (acquisition time 6:22) for STE-only analysis (Figure 5, bottom row), which amounts to an effective speedup of ∼4x purely through the use of both SE and STE in qT_1_ mapping. Furthermore, the precision tends to be more homogenous over the brain. The efficiency gain for high precision could be explained by the lower complexity of model fit performed in STE/SE ratio analysis, and the better homogeneity by a decreased confounding effect of T_2_, RF receive coil sensitivities and proton density on the fit. The efficiency gain is particularly interesting in the context of the theoretical efficiency of MESMERISED compared to e.g. IR-EPI. The use of a STE halves the signal intensity and SNR, which in principle requires an acquisition time factor of 4 to overcome and accelerating STEAM for qT_1_ mapping would therefore only seem to provide useful SNR efficiency benefits beyond a 4x speedup. However, these results indicate that effectively using both the half signal SE and half signal STE in STEAM can compensate for the 50% signal decrease, at least in terms of the precision of the resulting qT_1_ maps. Here it should be noted that good results for qT_1_ mapping are obtained at less than 100% duty cycle, with a doubling of the minimum achievable TR and TM (Figure 5, bottom row vs. top row), providing a better spacing and range of TM’s for T_1_ relaxometry. Potentially, in the future, this space can be used for simultaneous sampling of other contrasts, such as a multi-echo EPI readout (Poser and Norris, 2009) for qT_2_/qT_2_* mapping for further efficiency.

When compared to the MP2RAGE sequence (Marques et al., 2010), MESMERISED’s single-shot EPI readout strategy lends itself to sampling considerably more points on the full T_1_ decay curve (here 7 or 8 TMs vs. 2 inversion times for MP2RAGE) in less time, allowing a more straightforward relaxometry approach, but does so at a considerably lower resolution (here 1.8 to 1.25 mm vs. 1 to 0.6 mm for MP2RAGE). Similarly, the variable flip angle approach in DESPOT1 (Deoni, 2007) with spoiled gradient recalled echo (SPGR) images lends itself well for 1 mm resolution qT_1_ maps at 3 T, but contends with considerably sensitivity to B_1_^+^ inhomogeneities. In that sense MESMERISED’s T_1_ relaxation sampling strategy is closer to MS-IR-EPI (Panchuelo et al., 2021) and it could benefit from the multi-shot readout applied there to enable qT_1_ mapping with higher spatial resolution than presented here. The denser sampling of the decay curve in MESMERISED and MS-IR-EPI may lend itself to better characterisation of multi-component or multi-compartment qT_1_ mapping than MP2RAGE and DESPOT, particularly in the context of quantitative multi-parameter mapping and when spatially aligned diffusion MRI is needed.

### Diffusion MRI

For diffusion imaging, MESMERISED inherits from STEAM the capacity to achieve long diffusion times and high b-values, as well as the possibility to simultaneously generate diffusion-weighted SE images and diffusion-weighted STE images. MESMERISED adds to this a very high data rate to explore a large range of b-values, diffusion directions, or diffusion times in a short time. For instance, 370 whole-brain diffusion volumes over 8 b-value shells up to b = 7000 s/mm^2^ can be generated at 2 mm isotropic in 7 min and 47 s (ESxMB = 5×3, 2.95x speedup compared to MB-STEAM), a data rate of almost a volume per second, or at 1.8 mm isotropic in 10 min and 53 s (ESxMB = 6×3, 3.43x speedup compared to MB-STEAM), a data rate of more than a volume per 2 seconds. In addition, this is achieved with a lower diffusion gradient duty cycle (as well as less gradient coil heating and accompanying cooling requirements) than PGSE, since the longer diffusion times in STEAM dictate shorter diffusion gradient durations for the same maximum b-value. The shells consist of one half SE volumes in low-b shells which can support analysis with Gaussian diffusion models as in DTI, and one half STE volumes in the same number high-b shells which are suitable for non-Gaussian diffusion models, such as DKI, NODDI and CHARMED models. This provides a wealth of diffusion information from several different models in short acquisition time, which could be further extended in future work. For instance, instead of collecting a relatively large number of directions in 4 shells for each SE and STE, fewer (e.g. 6-12) directions could be acquired at each of many (e.g. 14-28) b-values. When carefully chosen, this would continue to support fitting of the aforementioned models on high-b STE volumes, while creating an opportunity to additionally fit the intra-voxel incoherent motion (IVIM) model (LeBihan, 1990) to the richly sampled low b-value domain in the SE volumes. Further tuning potential for the b-values of SE volumes (for a given STE max-b) lies in the choice of mixing time (and therefore diffusion time) of the STE volumes because, for the same TR and STE b-value, the SE b-value varies with different choices of TM and δ (see Supplementary Table 4).

### Comparison with MB PGSE for 7 T high-b multi-shell diffusion MRI

In a direct comparison for high-b multi-shell diffusion imaging at 7 T, MESMERISED displays markedly higher tSNR efficiency in white matter than MB PGSE at 7 T, when matched for RF-pulse regime, readout EPI train and image reconstruction. MESMERISED tSNR per unit time is ∼170% higher, and its tSNR efficiency is ∼70% higher in white matter than that of MB PGSE. The gains in gray matter are potentially even higher, but the comparison was aimed at white matter, since this is currently the main application domain of high-b diffusion MRI. To illustrate, this efficiency allows MESMERISED to acquire a 370 volume 8-shell diffusion dataset at 1.8 mm isotropic in ∼11 min, which would take MB PGSE ∼26.5 min to acquire at lower tSNR. The main factors in this are MESMERISED’s acquisition of both a high-b STE and low-b SE volume in a TR, and PGSE’s limitations of SAR and gradient duty cycle, which forces it to run at increased TR and TE. Of note here is that SAR and gradient limits each have a similar individual effect on PGSE TR: in the investigated case, SAR limits forced its TR upwards to 4 s and gradient limits forced it up a further ∼5% to 4.21 s. Therefore, avoiding only one of the limits would not shorten TR much to improve PGSE efficiency. MESMERISED avoids these limitations through a lower SAR and gradient load allowing full acquisition duty cycle. Combined with its effective use of both the diffusion weighted SE and STE, this allows it to overcome the 50% signal disadvantage (and the theoretical factor 4x acquisition time disadvantage) and outperform MB PGSE for 7 T high-b diffusion imaging.

B_1_^+^ maps at the employed reference voltage show that at 20% overflipping in the center of the brain, up to 50% underflipping occurred in other parts of the brain, despite the use of dielectric pads. This suggest that higher reference voltages could perhaps provide a higher overall level of tSNR over the brain, at the cost of lower tSNR in the center, higher SAR levels, and consequently increase in (SAR limited) TRs. This might affect PGSE more than MESMERISED since PGSE at MB = 2 has roughly 10% higher SAR per TR than MESMERISED at MB = 3. The SAR load for PGSE could be improved through parallel transmission (pTX) techniques and optimized MB pulses such as PINS (Norris et al., 2014; Norris et al., 2011), multi-PINS (Eichner et al., 2014) and pTx-MB (Poser et al., 2014; Wu et al., 2013) but the improved SAR would equally allow higher MB factors and higher efficiency in MESMERISED. The fact that B_1_^+^ inhomogeneity is one of the main drivers of spatial variations in tSNR and diffusion modeling uncertainty strongly suggests that methods for improved B_1_^+^ homogeneity, such as pTX or improved dielectric pad strategies, are desirable for both PGSE and MESMERISED.

Beyond efficiency, MESMERISED’s effectiveness in fiber orientation and microstructure modeling was qualitatively and quantitatively higher than that of PGSE. Diffusion modeling parameter maps (FS and FA) for both MESMERSED STE and SE volumes have excellent WM-GM contrast for several different echo-shift factors, whereas those for PGSE lack WM-GM contrast, particularly in deep GM nuclei, and appear to have an overall upward bias. Modeling precision quantified by uncertainty is markedly better for both MESMERISED STE and SE volumes than for PGSE, excluding DTI modeling at very low b-values. A significant factor beyond tSNR in the upward bias and higher uncertainty of FS for PGSE is the strong observed Gibbs ringing caused by the high water signal, likely through the long TE and strong resulting T_2_-weighting. The assessment of efficiency and effectiveness on 7 T multi-shell diffusion MRI was performed at b = 7000 s/mm^2^ with a high performing 70 mT/m gradient. The chosen b-value is non-arbitrary, as it has been shown that this is the minimum diffusion weighting necessary to fully suppress the contribution of extra-axonal water to the signal and obtain a high specificity to the intra-axonal compartment (more generally: the restricted compartment internal to thin axonal, glial or dendritic processes; Veraart et al., 2019), after the need for high b-values to accurately distinguish intra- and extra-cellular diffusion had already been made clear (Assaf and Basser, 2005; Assaf et al., 2004). Various types of 7 T PGSE-EPI acquisitions at moderate b-values (between ∼1000 and ∼2000 s/mm^2^) have yielded high quality data at very high spatial resolutions, highly suitable for axon orientation modeling and tractography (Eichner et al., 2014; Vu et al., 2015), including those with in-plane segmentation (Heidemann et al., 2010), and slice-dithered through-plane resolution enhancement (Setsompop et al., 2018). However, such acquisitions lack the intra-axonal signal specificity at high b-values, important for accurate diffusion modeling of axonal microstructure and integrity, which we targeted here. Therefore, we believe that while the domain of moderate-b high-resolution 7 T diffusion imaging may be best served by PGSE-EPI variants, MESMERISED is superior for high-b moderate resolution 7 T diffusion imaging. In the future, the limits imposed by gradient duty cycle for high-b PGSE could be improved by further gradient hardware developments beyond the employed generation of high performing 70-80 mT/m gradients. For PGSE, higher amplitudes, higher duty cycle, or both simultaneously, could strongly reduce its TE, which is dominated by diffusion gradient duration, while the already short TEs in MESMERISED might benefit less. On the other hand, low TE readout strategies, such as center-out spiral sampling, could strongly reduce TE in MESMERISED (which is dominated by readout train length), and provide proportionally more benefit than for PGSE.

### Varying diffusion time

The combination of high data rate and a wide range of achievable diffusion times makes MESMERISED an excellent tool to investigate the behavior of diffusion phenomena at long and varying diffusion times. Several studies have shown that processes such as exchange or mesoscopic structure due to random axonal packing leave their mark in the change of diffusion MRI measures with increasing diffusion time (Burcaw et al., 2015; Nilsson et al., 2013; Novikov et al., 2014). Diffusion time-resolved dMRI can even be used to achieve some sensitivity to axonal diameter distributions (Alexander et al., 2010; Assaf et al., 2008; De Santis et al., 2016b). Most of these applications require quite extensive sampling of diffusion times, b-values and, sometimes, directions, often leading to long (e.g. 30 - 60 min) acquisitions with current techniques. MESMERISED can achieve time-efficient sampling of different diffusion times in the STE volumes at otherwise matched parameters by increasing ES factor with increasing diffusion time. We demonstrate this with 1.8 mm isotropic diffusion data acquired at four diffusion times from ∼ 80 ms to ∼ 290 ms, which supports both DTI and DKI at each diffusion time (Figure 12). The analysis of the resulting STE data with DTI and DKI captures a dependence of mean diffusivity and mean kurtosis with increasing diffusion time in white matter (Figure 10D), in agreement with earlier reports (Fieremans et al., 2016; Jespersen et al., 2018). In the future, MESMERISED could be a useful tool in further investigating such micro- and meso-structural phenomena. In principle, less than the number of directions acquired here per b-value (24 for b = 1000 s/mm^2^, 48 for b = 2500 s/mm^2^ at a total acquisition time of ∼ 17 min for all diffusion times) might be sufficient for DKI and thus a shorter MESMERISED acquisition may be possible for this purpose. Furthermore, sampling higher diffusion times is a possibility, since correspondingly higher ES factors allow these to be acquired at the same TR (e.g. 430 ms at ES = 6 and 560 ms at ES = 8), and the performed PGSE comparison experiment shows that high tSNR is maintained with MESMERISED even at b-values up to b = 7000 s/mm^2^ and a diffusion time up to 430ms.

### Quantitative multi-contrast mapping

MESMERISED enables fast multi-contrast mapping over the whole-brain, for instance at 1.8 mm isotropic, with qT_1_ mapping, qT_2_ mapping, B_1_^+^ mapping and 370 volume multi-shell diffusion, in a little over 17 minutes. Accepting a lower precision level in the quantitative maps (i.e. working with a single, rather than three repetitions), and 200 volumes in the diffusion dataset (cf. Supplementary Tables 1-3), would lower the total multi-contrast acquisition time to under 10 minutes (i.e. qT1 mapping in 1:18, qT2 mapping 0:56, B_1_^+^ mapping 1:05 and 200 volume multi-shell diffusion in 6:15). Here, each modality includes ∼30 s of GRAPPA and MB reference scans, which in the future might be shared, providing another 1:30 time gain. Here we note again that qT_2_ and B_1_^+^ mapping, per se, do not benefit from the accelerating echo-shifting capacity in MESMERISED. They derive from basic properties of the STEAM sequence, i.e. relaxometry on SE or STE echo times and the α - 2α – α technique, respectively, which can be performed independently of any echo-shifting. However, the ability to perform qT_2_ and B_1_^+^ mapping with the same pulse sequence that provides highly accelerated qT_1_ and diffusion imaging has considerable advantages for fast multi-contrast mapping. First, all quantitative contrasts are generated by the same sequence and readout train, leading to similar image distortions which aid alignment of the different maps where needed. Second, for the same reason, the images from all contrasts share the same image noise distribution, which aids denoising with methods that take all images into account (Cordero-Grande et al., 2019; Moeller et al., 2021; Veraart et al., 2016). Third, well-aligned B_1_^+^ maps can help in correcting transmit field related biases in quantitative maps, for instance in qT_1_ maps (Marques et al., 2010). As also mentioned above, in comparison to quantitative multi-contrast mapping techniques based on a SPGR readout, such as DESPOT1/DESPOT2 (Deoni et al., 2005), MP2RAGEME (Caan et al., 2019) and MPM (Weiskopf et al., 2013), MESMERISED with EPI readout will more easily sample many more points on T_1_ and T_2_ decay curves, allowing a more straightforward relaxometry approach, but at a lower resolution. In its capacity for sampling qT_1_, qT_2_ and diffusion contrasts efficiently, MESMERISED is similar to the ZEBRA technique (Hutter et al., 2018), which is instead based on IR-prepared spin-echo EPI, and therefore more SAR intensive. This is less of a hindrance at 3T, and extensive sampling of the multi-dimensional [T_1_, T_2_*, b] contrast space could be achieved efficiently with ZEBRA, though at considerably lower spatial resolution (2.5 vs. 1.8 mm isotropic) and lower b-values (2600 vs. 7000 s/mm^2^) than those reported here. We believe the efficient sampling of the [T_1_, T_2_, B_1_, b, Δ] contrast space in MESMERISED is ideal for characterisation of multiple microstructural components or compartments, particularly when spatially aligned diffusion MRI with high b-values and/or varying diffusion times is needed.

### Outlook

As mentioned above, a useful extension will be a multi-echo EPI readout, which would enable further efficiency in mapping both qT_1_ and qT_2_/qT_2_* since unused duty cycle in optimal-TM sampling for T_1_ relaxometry can be used to sample multiple TE’s per shot for T_2_/T_2_* relaxometry. This could potentially also make use of efficient sparse sampling trajectories of the temporal relaxation curve, such as in echo-planar time-resolved imaging (EPTI; Wang et al., 2019). A multi-echo readout would also enable further speed-up of mixed or dependent contrast sampling, such as mixed T_2_/T_2_* relaxometry and diffusion imaging (Tax, 2017; Veraart et al., 2018), and help characterise tissue compartments with separate relaxation constants and diffusion behavior (De Santis et al., 2016a; Hutter et al., 2018; Wu, 2017). Future combination with SAR efficient MB pulses, such as PINS (Norris et al., 2014; Norris et al., 2011), multi-PINS (Eichner et al., 2014) and pTx-MB (Poser et al., 2014; Wu et al., 2013) could allow high data quality at potentially even higher MB factors. For the fastest acquisitions, acquisition of the reference data for GRAPPA and MB reconstruction takes a considerable proportion of the running time (about 30 s in the current implementation), which might be sped-up further with more efficient strategies. Additionally, a flexible table-based sequence looping implementation in which all parameters (TM, ES, TE, flip angles, diffusion direction and gradient amplitude) can be controlled independently, could allow running qT_1_, qT_2_, B_1_^+^ and diffusion mapping in a single sequence run, akin to magnetic resonance fingerprinting (Ma et al., 2013) in its Cartesian form (Benjamin et al., 2019). When EPI readouts are employed, shared reference scans and reversed phase-encode direction scans, would further decrease overhead time. Finally, the SAR efficiency of a STEAM-based sequence makes it very attractive for high duty cycle and multiband acceleration at 7 T, but many of the advantages could translate well to 3 T, particularly when MB factors are increased even further, as allowed by modern 32 to 64 channel head or head/neck RF-coils.

## Conclusion

MESMERISED achieves super-accelerated 7 T STEAM imaging by combining echo-shifting and multiband/simultaneous multi-slice acceleration. This leads to very high multiplicative acceleration factors and time efficiency for qT_1_ and diffusion imaging. MESMERISED can probe combined T_1_, T_2_ and diffusion contrast with high time efficiency for fast multi-contrast mapping and characterization of multi-component relaxometry, diffusion, and exchange.

## Acknowledgements

AR and FJ were supported by an ERC Starting Grant (MULTICONNECT, #639938). AR was further supported by a Dutch science foundation (NWO) VIDI Grant (#14637). FJ was partially supported by the German Research Foundation (DFG Emmy Noether Program: MO 2397/4-1). BP is partially funded by NWO VIDI Grant 016.Vidi.178.052. and NIH R01 MH111444/MH/NIMH.

