## Supplementary Information for "MESMERISED: Super-accelerating T_1_ relaxometry and diffusion MRI with STEAM at 7 T for quantitative multi-contrast and diffusion imaging"

### Supplementary Tables

| ***Resolution - slices*** | ***MB - GRAPPA*** | ***TR [s] / TE [ms]*** | ***ES (TM[ms])*** | ***ESxMB*** | ***Acquisition time 1/3/5 reps [m:s]*** |
|---|---|---|---|---|---|
| 1.8 mm – 72 slices | 3 - 2 | 6.90 / 28 | 1(140), 2(280), 3(420), 4(560), 6(840), 8(1120), 12(1680) | 3, 6, 9, 12, 18, 24, 36 | 1:18 / 2:07 / 4:32 |
| 1.25 mm – 108 slices | 3 – 3 | 4.4 / 31.70 | 1(60), 2(120), 3(180), 4(240), 6(360), 9(540), 12(720), 18(1080) | 3, 6, 9, 12, 18, 27, 36, 54 | 1:05 / 2:16 / 3:26 |
| 1.25 mm – 108 slices | 3 – 3 | 8.8 / 31.70 | 1(120), 2(240), 3(360), 4(480), 6(720), 9(1080), 12(1440), 18(2160) | 3, 6, 9, 12, 18, 27, 36, 54 | 1:40 / 4:01 / 6:22 |

**Supplementary Table 1.** Quantitative $T_1$ imaging parameters. Total acquisition times are for 1, 3 and 5 repetitions for each ES/TM and including 30 seconds of GRAPPA and MB reference scans. All $qT_1$ imaging uses standard Exc-Sto-Rec = 90°-90°-90° flip angles.

| ***Resolution - slices*** | ***MB - GRAPPA*** | ***TR [s] / TE [ms]*** | ***ES (TM[ms])*** | ***TE[ms] or Flip angle $\alpha_{TX}$[deg]*** | ***Acquisition time 1/3/5 reps [m:s]*** |
|---|---|---|---|---|---|
| 1.8 mm – 72 slices | 3 - 2 | 5.2 / variable | 1 (70) | TE = 28.3, 38.3, 48.3, 68.3, 88.3 | 0:56 / 1:48 / 2:40 |
| 1.8 mm – 72 slices | 3 - 2 | 5.2 / variable | 3(420) | TE = 28.3, 38.3, 48.3, 68.3, 88.3 | 0:56 / 1:48 / 2:40 |
| 1.8 mm – 72 slices | 3 - 2 | 6.9 / 28.3 | 1 (140) | $\alpha_{TX}$ = *30, 50, 70, 90, 110* | 1:05 / 2:14 / 03:23 |

**Supplementary Table 2.** Quantitative $T_2$ and $B_1$+ imaging parameters. $T_2$ imaging (row 1 and 2) uses standard Exc-Sto-Rec = 90°-90°-90° and no further corrections, e.g. for imperfect slice profile, were applied. B1+ imaging (row 3) uses flip angle combinations of Exc-Sto-Rec = α°- 2α°- α° for different multipliers α. Total acquisition times are for 1, 3 and 5 repetitions for each TE or flip angle combination and including 30 seconds of GRAPPA and MB reference scans.

| *Resolution - slices* | *MB - GRAPPA* | *TR [s] / TE [ms]* | *ES (TM[ms])* | *b[s/mm²] (#dir) STE echo* | *Acquisition time- Nr. vol - s/vol* |
|---|---|---|---|---|---|
| 2.0 mm – 60 slices | 3 - 2 | 2.36 / 40.4 | 5(290) | b0 (17), 1750 (24), 3500 (36), 5250 (48), 7000 (60) | 7:47 370 vols – 1.18 s/vol |
| 1.8 mm – 72 slices | 3 - 2 | 3.45 / 49.8 | 2(140) | b0 (17), 1750 (24), 3500 (36), 5250 (48), 7000 (60) | 11:08 370 vols – 1.73 s/vol |
| 1.8 mm – 72 slices | 3 - 2 | 3.41 / 46.0 | 3(210) | b0 (17), 1750 (24), 3500 (36), 5250 (48), 7000 (60) | 11:08 370 vols – 1.73 s/vol |
| 1.8 mm – 72 slices | 3 - 2 | 3.37 / 40.8 | 6(420) | b0 (17), 1750 (24), 3500 (36), 5250 (48), 7000 (60) | 10:53 370 vols – 1.69 s/vol |
| 1.8 mm – 72 slices | 3 - 2 | 3.40 / 49.9, 3.37 / 46.2, 3.36 / 43.8, 3.35 / 40.9 | 2(140), 3(210), 4(280), 6(420) | b0 (6), 7000 (60) + b0 (27) | 4:14 + 2:02 4:12 + 2:01 4:12 + 2:01 4:11 + 2:01 1.7 - 1.675 s/vol |
| PGSE 1.8 mm – 72 slices | 2 - 2 | 4.21 / 97.8 | n.a. | b0 (6), 7000 (60) + b0 (27) | 5:08 + 2:24 4.21s/vol |
| 1.8 mm – 72 slices | 3 - 2 | 3.50 / 45.0 | 1(70), 2(140), 3(210), 4(280) | b0 (8), 1000 (24) 2500 (48) | 16:50 560 vols – 1.75 s/vol |

**Supplementary Table 3.** Diffusion imaging parameters. Total acquisition times are for all combinations of ES/TM and b-values and including 30 seconds of GRAPPA and MB reference scans. Number of volumes in the final column is SE and STE volumes for all b-shells and ES/TM. B-values are listed for the STE echo volume.

| **ES** | **TM** | **δ** | **b-val SE 4/4, 3/4, 2/4, 1/4** |
|---|---|---|---|
| 6 | 420 ms | 7.37 ms | 223, 167, 112, 56 s/mm² |
| 4 | 280 ms. | 8.65 ms | 350, 264, 176, 88 s/mm² |
| 3 | 210 ms | 9.83 ms | 483, 362, 241, 120 s/mm² |
| 2 | 140 ms | 11.71 ms | 752, 564, 376, 188 s/mm² |

**Supplementary Table 4**. Combinations of echo-shift factor (ES), mixing time (TM), diffusion gradient length (δ), and Spin-echo b-value achievable at the same TR for the case of 1.8 mm isotropic, maximum STE b-value = 7000 s/mm2, at full duty cycle for GRAPPA factor 2 and multiband factor 3. Here $\Delta_{SE} = \delta + 8.88$ ms, $\Delta_{STE} = TM + \Delta_{SE}$ and $|G| = 70$ mT/m. The listed SE b-values are for a 4-shell acquisition with equally distributed b-values (i.e. at 4/4, 3/4, 2/4 and 1/4 of max-b), corresponding to STE b-value = 7000, 5250, 3500, 1750 s/mm$^2$. Note that full-duty cycle values also depend on EPI readout train length, and therefore values would be slightly different for different resolutions and GRAPPA factors.

### Supplementary Figures

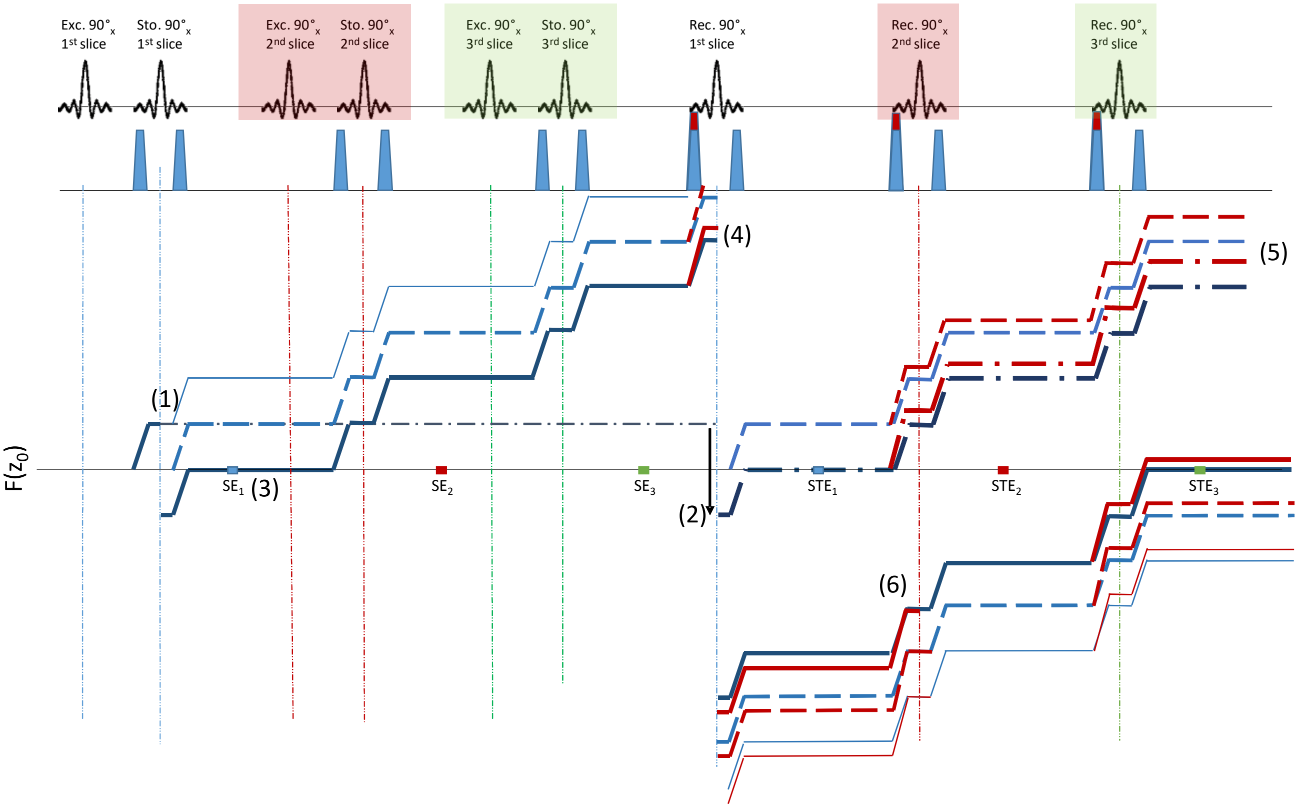


**Supplementary Figure 1**: Extended phase diagram (EPG) for the diffusion/crusher gradients around storing and recalling pulse of MESMERISED for the case of ESxMB = 3x1 (i.e. no MB, single slice). Here the 1st, 2nd and 3rd free induction decay (FID) pathways from, respectively, the excitation pulse (solid line with different thickness), storing pulse and recalling pulse (dashed lines) pulses for the first slice are shown, resulting in (all) the echo pathways of SE and STE block of the first slice, including first, second, third and fourth SE, plus the STE pathway. Here, the pulses for further slices (in red and green) are assumed not to interact with these pathways, and influence of other gradients (including slice select gradients and readout EPI train) are neglected. The 1st FID pathway generates the resulting first spin-echo (SE, at 3) after the storing pulse refocuses the FID (at 1), and the stimulated echo (STE) signal when the stored magnetization (dash-dotted line at 1) is recalled by the corresponding recalling pulse (2). The extensions of the 1st FIDs in case of imperfect storing or rephasing of the Storing and Recalling pulses are also shown. Formation of the second and third SE is avoided by the crusher configuration. A possibility for contamination is the 4th SE, from the 1st FID pathway, where after generating the primary SE (at 3), the pathway (the thicker solid line from the storing pulse at 4) could contaminate the STE from slice 3 (at 7). However, this pathway undergoes $T_2$ decay for TE + ~1.7 TM before potentially interacting with the later STE, resulting in extensive decay before contamination, except in the case of a long $T_2$ and very short TM. Additionally, here it is shown how an imbalanced 3rd diffusion/crusher gradient moment (red increment on top of the gradient) increases the dephasing in the pathways (red lines) and removes any contamination of the residual echo pathways) with the later STEs (7). This effect is enhanced for the 3rd FID, where this imbalanced gradient is more frequently applied (5). However, this extra dephasing is “reversed” when an equal number of imbalanced diffusion gradients is applied after the emission of the recalling pulse (6).

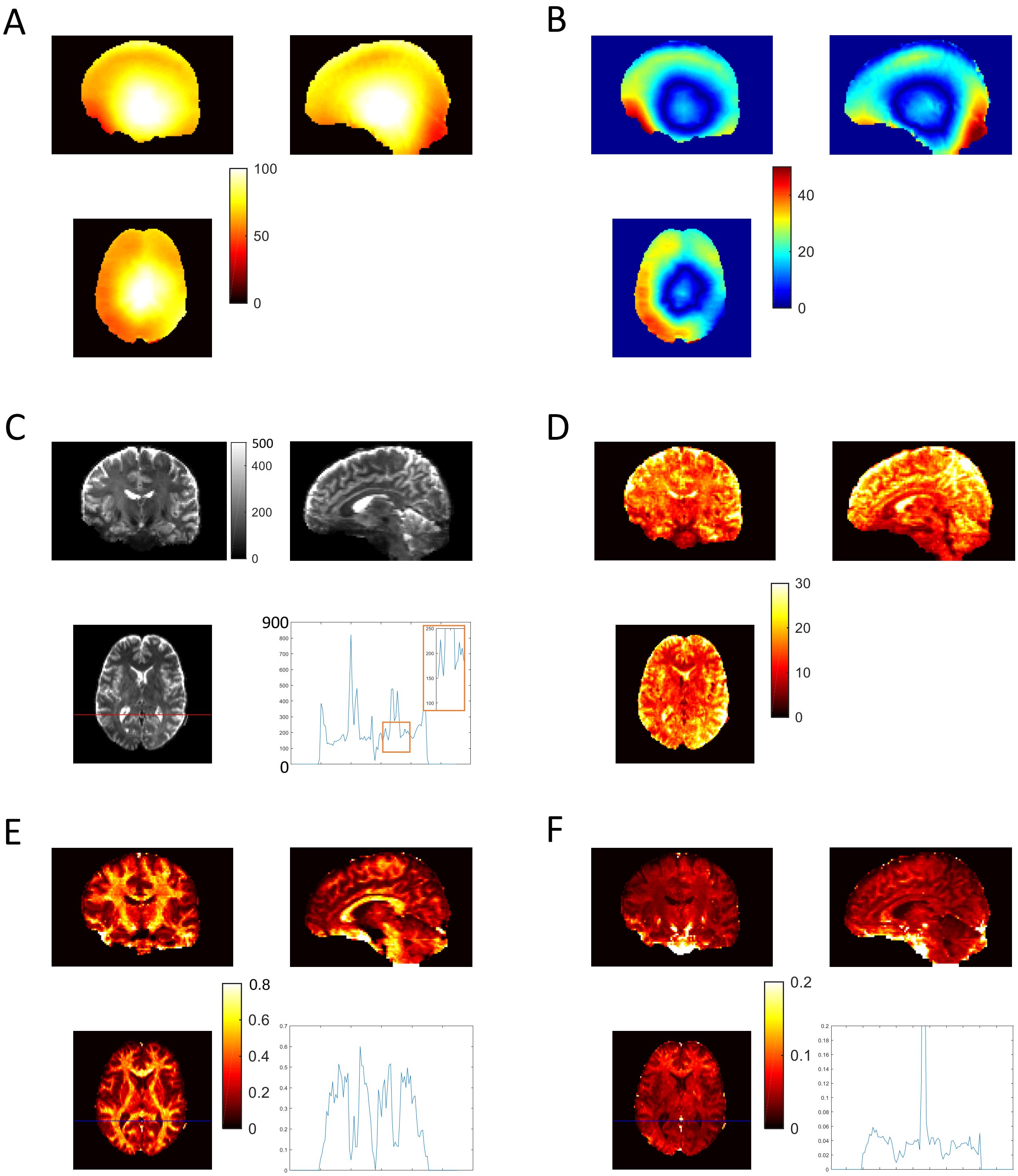


**Supplementary Figure 2**. Three orthogonal plane results for MESMERISED MB3 – ES3 STE volumes (c.f. Figure 9B, top) including $B_1^+$ maps and line plots. (A) Absolute $B_1$+ map in units of degrees (°) (B) the deviation of the local flip angle from the nominal 90° for the $B_1$+ map. (C) The mean $b_0$ volume for high-b multi-shell diffusion imaging, with a line plot over the red line in the transverse plane (inset: zoom around the ventricle) (D) tSNR map over $b_0$ volumes (E) The Fraction of Sticks (FS) map and (F) its uncertainty for high-b diffusion and $b_0$ volumes, with a line plot over the blue line in the transverse plane.

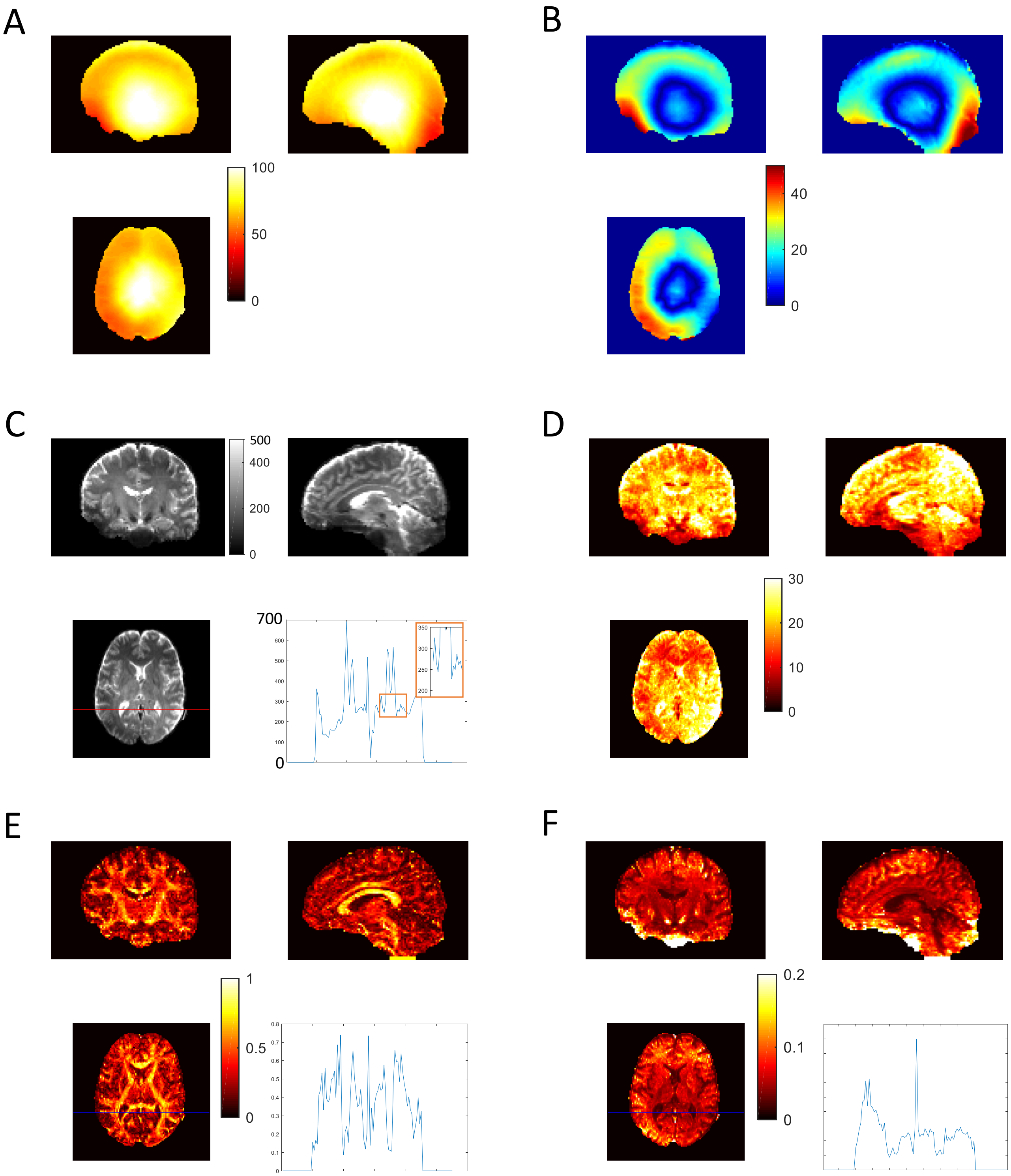


**Supplementary Figure 3**. Three orthogonal plane results for MESMERISED MB3 – ES3 SE volumes (c.f. Figure 9B, bottom) including $B_1$ maps and line plots. (A) Absolute $B_1$+ map in units of degrees (°) (B) the deviation of the local flip angle from the nominal 90° for the $B_1$+ map. (C) The mean $b_0$ volume for high-b multi-shell diffusion imaging, with a line plot over the red line in the transverse plane (inset: zoom around the ventricle) (D) tSNR map over $b_0$ volumes (E) The Fraction of Sticks (FS) map and (F) its uncertainty for high-b diffusion and $b_0$ volumes, with a line plot over the blue line in the transverse plane.

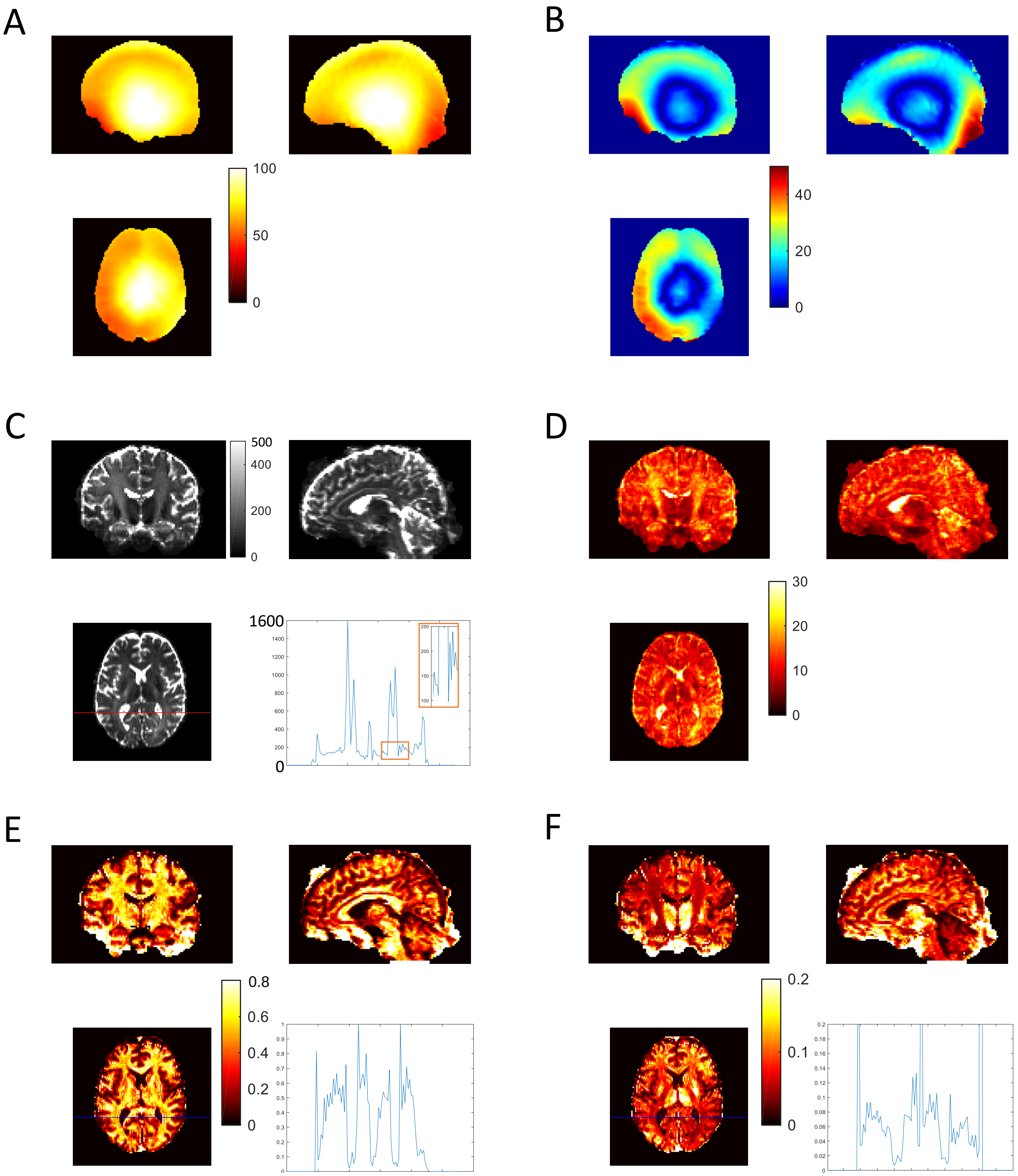


**Supplementary Figure 4**. Three orthogonal plane results for PGSE MB2 volumes (c.f. Figure 9E) including $B_1$+ maps and line plots. (A) Absolute $B_1$+ map in units of degrees (B) the deviation of the local flip angle from the nominal 90° for the $B_1$+ map. (C) The mean $b_0$ volume for high-b multi-shell diffusion imaging, with a line plot over the red line in the transverse plane (inset: zoom around the ventricle) (D) tSNR map over $b_0$ volumes (E) The Fraction of Sticks (FS) map and (F) its uncertainty for high-b diffusion and $b_0$ volumes, with a line plot over the blue line in the transverse plane.

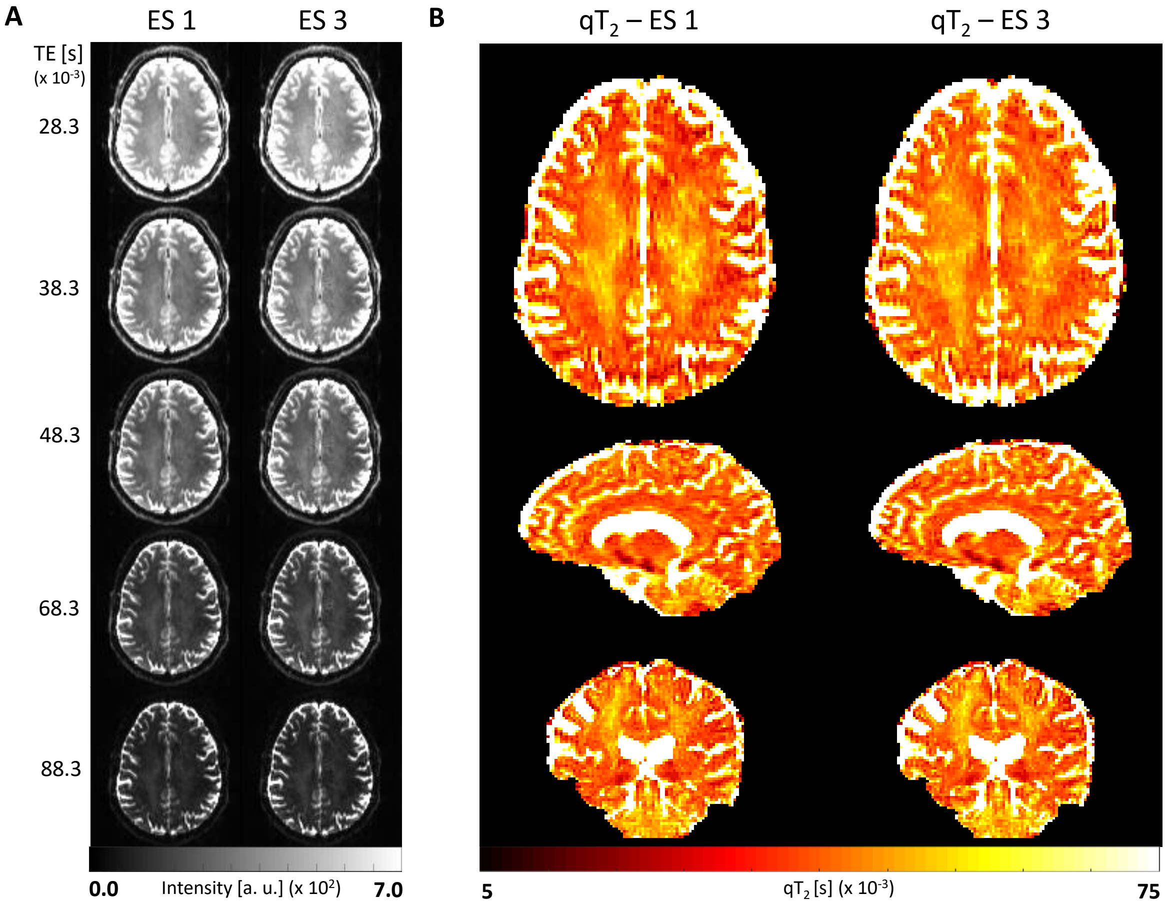


**Supplementary Figure 5:** MESMERISED $qT_2$ mapping and $T_2$w relaxometry data at 1.8 mm isotropic resolution. (A) Acquired SE volumes over echo times of 28.3 ms to 88.3 ms (see Supplementary Table 2) illustrating $T_2$ decay, qualitatively similar for ES 1 and ES 3 for both ES = 1 (left) and ES = 3 (right). (B) The corresponding $qT_2$ maps for ES = 1 (left) and ES = 3 (right); the $qT_2$ map seems affected by the echo-shifting mechanism in MESMERISED.

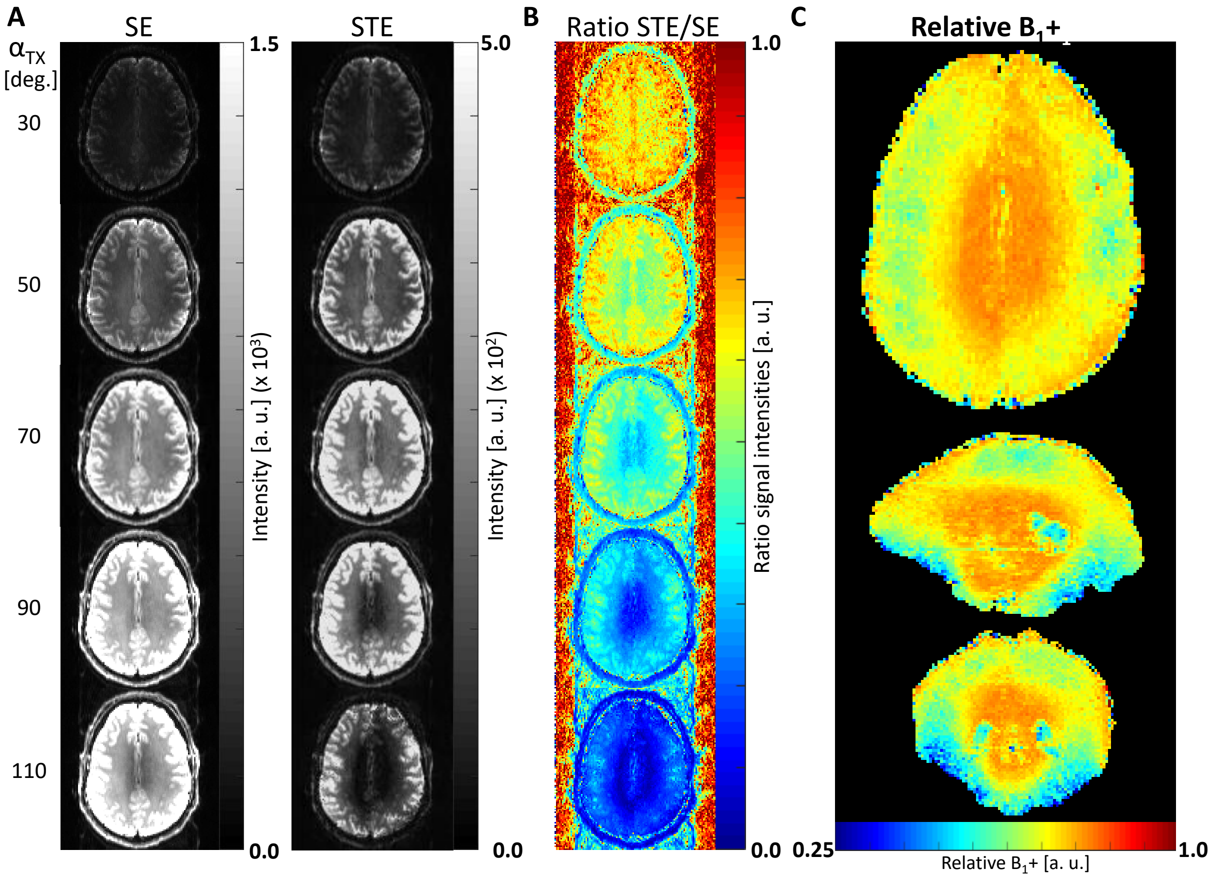


**Supplementary Figure 6**: MESMERISED B1+ mapping at 1.8 mm isotropic, derived from both the SE and STE volumes for different $\alpha_{TX}$ multipliers in an $\alpha_{TX}$ - $2\alpha_{TX}$ - $\alpha_{TX}$ flip angle scheme for Exc – Sto – Rec pulses. (A) Acquired SE (left) and STE (right) volumes for different $\alpha_{TX}$ multipliers (B) The corresponding SE/STE ratios used for $B_1$+ map fitting (C) The resulting whole-brain relative $B_1$+ map.
